# Transfer learning reveals species-specific olfactory preferences in Diptera and informs pest management strategies

**DOI:** 10.64898/2026.07.03.736362

**Authors:** Yifeng Zhang, Li Xu, Chenxi Gao, Tianmin Zhang, Suyang Duan, Yangming Yin, Xuanxiao Yang, Qingbin Sun, Xinghua Qin, Gang Li, Chang Xu, Hongbo Jiang, Huimeng Lu

## Abstract

The interaction between volatile organic compounds (VOCs) and odorant receptors (ORs), termed VOI, constitutes the molecular basis for species to recognize chemical information in the environment. Characterization of subtle differences in VOI relationships among closely related species is crucial for understanding the rapid evolution of ORs and for identifying species-specific chemical cues relevant to pest management. However, progress in this field has been constrained by both the scarcity of experimental data and limitations in computational prediction accuracy. To address this, we developed a virtual screening-enhanced transfer learning strategy that integrates large-scale molecular docking data with sparse functional experimental data. The resulting VOI prediction model was validated through functional experiments on the pest species *Bactrocera dorsalis*, demonstrating its cross-species predictive capacity. Using this model, we investigated the relationship between ecological niches and olfactory sensitivity in Diptera insects from multiple perspectives, with an emphasis on patterns that may inform pest behavioral research. The model’s predictive reliability was further supported by its consistency with known olfactory trends: it recapitulated the preferential responses of fruit flies and mosquitoes to esters and aromatics, respectively, and reproduced the previously reported negative correlation between olfactory and visual capacities in Drosophila. As a proof-of-concept application, we compared hematophagous and non-hematophagous mosquitoes, revealing overlapping chemical spaces but distinct patterns in their specifically recognized compounds. This study provides a reliable VOI prediction framework that reveals species-level olfactory preference patterns in Diptera, offering a computational pipeline to accelerate the discovery of behaviorally active compounds for species-specific pest monitoring and control.

## 1 Introduction

The volatile organic compounds (VOCs) and odorant receptors (ORs) interaction (VOIs) serves as the molecular basis for insects to perceive complex chemical information in their environment(1). Insects utilize dozens to hundreds of ORs to detect tens of thousands of VOCs in their surroundings, contributing to their strong environmental adaptability and complex social behaviors. The function of ORs is determined by the activation response elicited upon its binding to specific VOCs. During OR evolution, gene duplication, functional differentiation, pseudogenization and loss have enabled the gradual renewal of OR repertoires and the emergence of new functions(2–5), collectively shaping the functional diversity of ORs across different species. The functional characteristics of insect OR repertoire largely determine its olfactory potential. For closely related species, quantifying and comparing the functional divergence of their OR repertoires is crucial for understanding the molecular mechanisms of olfactory adaptation. This approach facilitates an understanding of the intrinsic links between OR functional evolution and environmental adaptation, providing insights into how natural selection shapes olfactory potential to accommodate diverse ecological niches, and may help identify olfactory features relevant to pest behavioral research.

The experimental method for functional characterization of ORs mainly involves constructing *in vivo* or *in vitro* protein expression systems for physiological and biochemical testing to determine the binding strength of VOIs. Different insect odorant receptor expression systems have been constructed(6–10), and a number of studies have reported the functional characterization of insect ORs(6, 7, 10–12). However, due to experimental difficulties and time constraints, the target species are often limited to common insect species, such as fruit flies(1, 7), mosquitoes(9, 13), moths(14, 15), honeybees(16), etc., and the number of tested odor molecules is also very limited. The functions of the large diversity of ORs among multiple insect species remain unknown. Additionally, due to the significant divergence in ORs among different insect species(17, 18), achieving cross-species VOI prediction based on existing experimental data remains a different problem. Unevenly distributed experimental data pose a challenge for constructing species-level OR functional profiles, thereby hindering systematic comparisons of olfactory potential among closely related species and limiting the discovery of behaviorally relevant compounds for pest species. Using computational methods to predict VOI relationships can accelerate the functional characterization of ORs and enable the investigation of olfactory evolution at broader scales of species and chemistry. Therefore, it is imperative to explore effective methods for predicting VOIs that can facilitate large-scale functional profiling and candidate prioritization for behavioral studies.

In recent years, significant progress has been made in the structural analysis of insect ORs(19–22), which helps us further understand the molecular mechanisms of VOI and conduct more accurate VOI analysis. Although the number of insect OR structures identified by experimental methods is limited, they can be extrapolated through recently developed protein structure modeling techniques(23, 24) to predict the structure of OR protein structures. Therefore, we can also predict protein functions by molecular docking. Molecular docking involves searching for interactions between ligands and receptors and determining the optimal binding mode in the region of the receptor active site through the principle of spatial structure complementarity and energy minimization. Similar to the role of virtual screening in drug discovery, molecular docking can enrich potential ligands with binding ability from a large number of VOCs(25), improving the efficiency of OR functional characterization(26, 27), while also providing an initial overview of species-level VOI binding trends relevant to olfactory ecology(28, 29). On this basis, we previously developed a virtual screening database called iORbase(30). Compared to experimental methods, virtual screening can provide rich data to reveal the binding trends of ORs and ligands, thus compensating for the lack of experimental data. However, the limited precision and false-positive rate of docking hinder its ability to resolve ecological differences. This renders it inadequate for comparative studies of specific adaptive traits, such as discerning olfactory divergence between closely related species, where higher-resolution VOI predictions are indispensable. Furthermore, while molecular dynamics (MD) simulations can analyze structure-activity relationships to improve VOI prediction accuracy(26, 31, 32), they require substantial computational resources, making them impractical for high-throughput applications. These limitations highlight the need for a prediction strategy that balances accuracy with scalability across large chemical spaces.

Previous studies using conventional machine learning methods have primarily focused on predicting potential agonists for target ORs based on VOC characteristics(33–36), or forecasting responses of unknown ORs to target VOCs through sequence similarity analysis(37). While these approaches have achieved some success, they demonstrate limited efficacy when dealing with broader and more complex VOI relationships. The deep learning method provides more powerful models that can be used to predict protein-molecule interactions(38–42). This method also has great potential in identifying the function of ORs. Based on the experimental OR data, deep learning models can be constructed to achieve rapid prediction of the VOI. However, the establishment of deep learning models requires a large amount of data, and limited existing experimental data make it difficult to meet this demand. Since the available experimental data are limited to a narrow range of ORs with high sequence variability, training a robust model with these data alone is challenging. Transfer learning offers a promising solution by allowing a model pretrained on large-scale data to be adapted to data-scarce tasks, as demonstrated in previous studies (43–47).

To address the data scarcity in VOI prediction, we developed a transfer learning strategy that integrates the broad coverage of molecular docking with the precision of experimental data. Based on this strategy, we developed insectOlf, a framework designed to reliably quantify and compare olfactory potential across species. We designed and conducted functional experiments on *Bactrocera dorsalis* to validate its predictive capability. Using this framework, we assessed its predictive reliability by testing its consistency with known olfactory trends across Diptera, and performed a proof-of-concept exploration by comparing hematophagous and non-hematophagous mosquitoes. In conclusion, this work establishes a computational approach for VOI profiling that combines docking-based pretraining with experimental fine-tuning. The framework, validated through both cross-validation and functional experiments, enables cross-species olfactory comparison and may facilitate the prioritization of candidate compounds for pest behavioral research. We anticipate that this framework will serve as a valuable tool for developing targeted semiochemical-based strategies in integrated pest management (IPM).

## 2 Method and Materials

### 2.1 Filtering and processing of the functional datasets

To construct the OR function prediction model, we selected the OR functional characterization results of four species, *Drosophila melanogaster* (*D. melanogaster* dataset)^4^, *Anopheles gambiae* (*A. gambiae* dataset)(6), *Locusta migratoria* (*L. migratoria* dataset)(11) and *Spodoptera littoralis* (*S. littoralis* dataset)(14). In addition, we selected high-volume molecular docking data from iORbase(30), containing approximately 14,000,000 pairs of interactions for 5980 ORs of 59 species and 2077 VOCs, for pretraining the model.

For the molecular docking data used for pretraining, the interactions between ORs and small molecules were described by the affinity score; the value of the affinity score was meaningful when it was negative, and a larger absolute value of the affinity score represented stronger potential binding between the corresponding protein and small molecule. For the molecular docking results of vina, a value less than 0 indicates a binding relationship between OR and VOC, and the larger the absolute value of the result is, the stronger the binding relationship is, while a value greater than 0 is meaningless (1.69% of the total data); therefore, we classified a value greater than 0 as 0 and took the remaining results as absolute values to ensure that the molecular docking results were consistent with the overall trend of the experimental results.

Due to variations in the data distribution ranges across different datasets, all data were preprocessed and scaled to the range of 0 to 1.

### 2.2 Molecular docking between ORs and VOCs

We utilized AutoDock Vina (v.1.2.0) (48) to perform molecular docking of ORs and VOCs. The 3D structures used for docking, namely, the OR PDB format file and VOC SDF file, obtained from iORbase(30) and PubChem(49), respectively, were converted to PDBQT format using Raccoon (v.1.0) software(50) to facilitate subsequent docking analysis. Prior to employing vina for molecular docking, it is crucial to determine the protein’s binding pocket in advance. To accurately identify the binding pocket of OR, we selected OR (PDB ID: 7LID) with a ligand structure as our template. After aligning the target protein with the template, a region centered on the template ligand within a radius of 20 Å was chosen as the binding pocket for our target protein. Ultimately, we extracted the top-ranking binding free energy from the resulting file in PDBQT format as our final outcome for molecular docking.

### 2.3 Model construction for deep learning

We used the Python package PyTorch (v. 1.10.2)(51) to quickly build our model and Scikit-learn (v.1.0.2) (52) and Numpy (v.1.21.5) (53) to process the data. We input the sequences of proteins and SMILES sequences of small molecules into the feature extractor for proteins and small molecules, respectively, to obtain the feature sequences of proteins and small molecules. The feature sequences of proteins and small molecules were merged as inputs to the neural network, and the corresponding affinities of proteins and small molecules were used as labels. The MSELoss function in PyTorch was used to measure the difference between the affinity values predicted by the model and those in the original dataset. The model adjusted the parameters according to the differences during training, and when the differences tended to stabilize, the model converged.

The model was pre-trained using all molecular docking data from iORbase, followed by fine-tuning on real experimental data. Specifically, we saved the model parameters obtained from training on the molecular docking data and loaded these parameters (rather than random initialization) before training the model on the real experimental data. Subsequently, a full fine-tuning strategy was adopted to update all layers of the model. The same hyperparameters were used for both the pre-training and fine-tuning stages.

For the real experimental data, we used 5[fold cross[validation to split the training and test sets, in order to evaluate the model’s performance on different datasets. For datasets containing multiple species (e.g., AgDm and Mix), we applied stratified sampling to ensure that the proportions of different species were consistent between the training and test sets.

### 2.4 Characterization of the OR function of *Bactrocera dorsalis*

The binding ability of candidate ORs to VOCs was determined by two-electrode voltage clamp electrophysiological recording assays as previously reported(54). Briefly, the verified PCR products of candidate *B. dorsalis* OR genes and BdorOrco were subcloned into a modified pT7Ts vector based on previous study (55). The recombinant plasmids were extracted using the plasmid MIDIprep kit (QIAgen, Düsseldorf, Germany) and the cRNA was synthesized using linearized plasmid with mMESSAGE mMACHINE T7 Kit (Invitrogen, Thermo Fisher Scientific, Lithuania) following manufacturer’s instructions. The purified cRNA was diluted to 2 µg/µL with nuclease-free water and stored at −80◦C. The oocytes from *Xenopus laevis* were dissected and separated using 1.5 mg/mL collagenase[(GIBCO, Carlsbad, CA, USA) in washing buffer (96 mM NaCl, 5 mM MgCl2, 2 mM KCl, and 5 mM HEPES [pH = 7.6]). The cRNA of BdorORs and BdorOrco were co-injected into selected healthy oocytes and incubation at 18◦C for 2 days in 1 × Ringer’s solution (96 mM NaCl, 2 mM KCl, 5 mMMgCl2, 0.8 mM CaCl2, and 5 mM HEPES [pH 7.6]) supplemented with 5% dialyzed horse serum, 50 mg/mL tetracycline, 100 mg/mL streptomycin, and 550 mg/mL sodium pyruvate. The VOC (≥98%, Sigma-Aldrich, USA) stock solutions (1 M) were prepared in DMSO (Sigma-Aldrich, USA) and diluted to indicated concentration with 1 × Ringer’s solution. The currents of injected oocytes induced by VOCs were recorded by two-electrode voltage clamp using OC-725C amplifier (Warner Instruments, Hamden, CT, USA). The nuclease-free water injected cells were served as negative control. The holding potential and the low-pass filter were −80 mV and 50 Hz, respectively. Digidata 1550A (Warner Instruments, Hamden, CT, USA) and pCLAMP10.5 software (Axon Instruments Inc., Union City, CA, USA) were used to acquired and analyzed the data, respectively. Data analysis and concentration-dependence curve were conducted with GraphPad Prism v8.0 software.

## 3 Results

### 3.1 Transfer learning enhances the generalization performance of the VOI predictor

Reliable prediction of the VOI relationships between OR repertoires and VOCs serve as the foundation for further analysis of insect olfactory ability. However, the scale of existing functional experimental datasets is extremely limited, failing to provide sufficient feature information for deep learning models, this limitation restricts the generalization performance of machine learning models across large-scale chemical spaces.

#### 3.1.1 Transfer learning strategy based on molecular docking

By integrating the advantages of massive molecular docking datasets and experimental datasets, a transfer learning framework was established (Fig. 1a). First, the model was pretrained on large-scale molecular docking data to enable it to learn the global feature distribution associated with VOI relationships. Subsequently, the pretrained model was fine-tuned using limited experimental data to adapt it to the target experimental dataset, this constitutes the transfer learning process. Through iterative optimization with experimental data, the model can derive a more precise VOI distribution pattern. Finally, leveraging the model pretrained on docking data and optimized by experimental data, zero-shot predictions can be performed for insects with unknown VOI relationships. The reliability of these predictions will be validated in the subsequent sections.

**Fig. 1.**
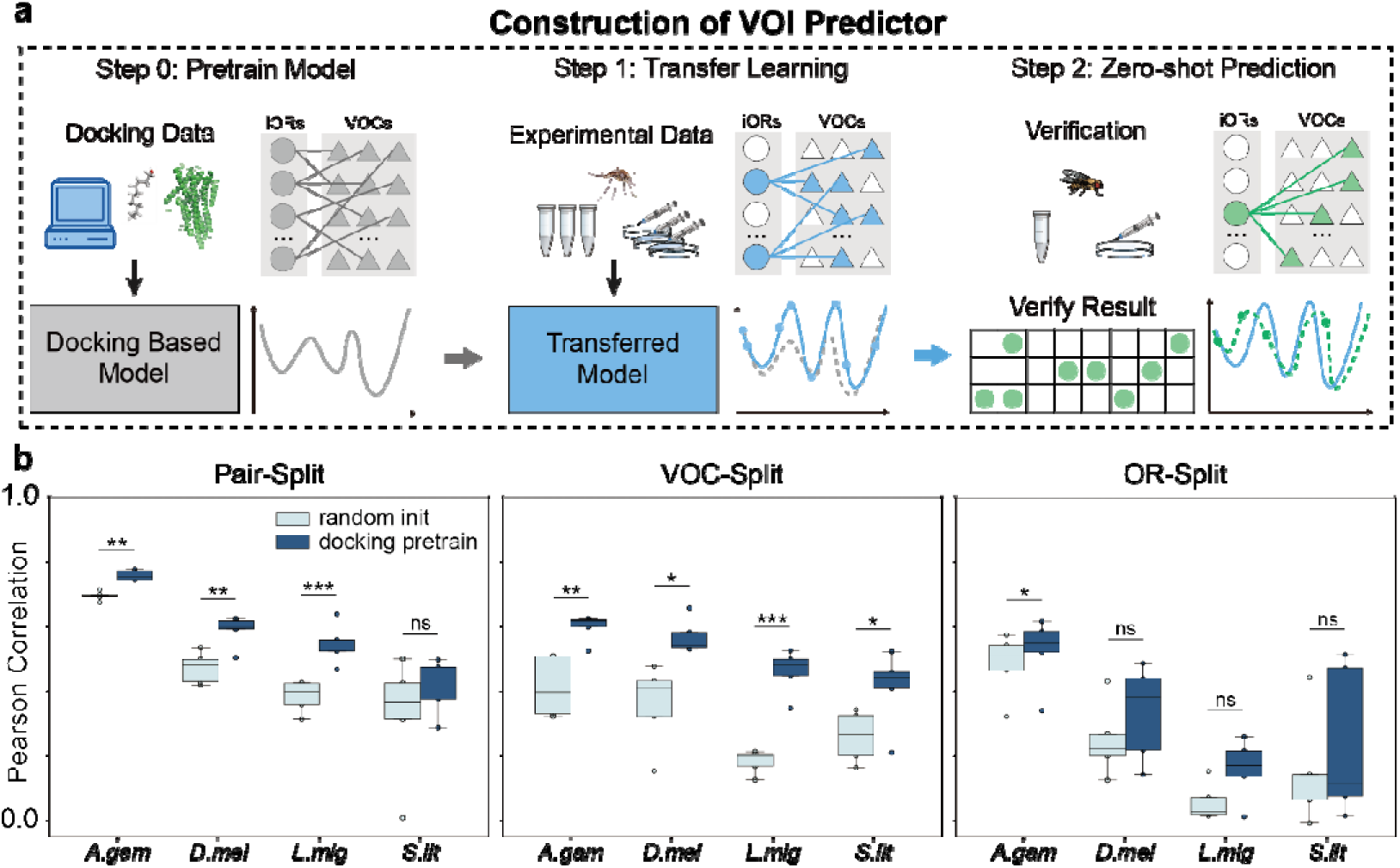
Transfer learning enhances the generalization performance of the VOI predictor. **a**, Schematic of the transfer learning strategy: pretraining on molecular docking data followed by fine-tuning on experimental data. **b,** Performance comparison between docking-pretrained and randomly initialized models across different experimental datasets.

**Tab. 1:** Dataset details.

| Dataset | ORs | VOCs | Sample Pairs |
| --- | --- | --- | --- |
| *Drosophila melanogaster* | 24 | 107 | 2568 |
| *Anopheles gambiae* | 50 | 100 | 5000 |
| *Spodoptera littoralis* | 17 | 35 | 595 |
| *Locusta migratoria* | 41 | 190 | 7790 |
| iORbase Docking | 5980 | 2077 | 14,000,000 |

#### 3.1.2 The cold start problem of the VOI predictor

To more accurately evaluate the ability of insects to recognize complex volatile organic compounds (VOCs) in the environment, our VOI predictor requires strong generalization ability to VOCs. In other words, we aim to investigate the model’s performance when confronted with unseen VOCs, which refers to the cold-start problem of the model. We employed two distinct splitting strategies to partition the training and testing sets. For the Pair split, we evaluated model generalization using three data-split strategies: (1) random split (in-distribution, ID), where training and test sets share ORs and VOCs; (2) VOC-split (out-of-distribution, OOD), where test VOCs are completely absent from training; and (3) OR-split (OOD), where test ORs are unseen during training. For each strategy, we performed 5-fold cross-validation on the four species datasets. As shown in Fig. 1b, transfer learning significantly improved prediction under both ID (Pair-split) and OOD (VOC-split) settings (paired t-test, p-values ranged from <0.001 to <0.05), achieving Pearson correlation coefficients above 0.5 for most datasets and up to 0.8 in the best case. However, performance on OR-split remained limited, highlighting the challenge of generalizing to completely novel receptors. These results suggest that the VOI predictor generalizes reasonably to unseen VOCs.

### 3.2 Predictive ability of the VOI predictor for different insect species

In the preceding sections, we have demonstrated the predictive ability of the VOI predictor for VOCs. Beyond this, we aim for the VOI predictor to effectively evaluate the olfactory ability of different insect species. Therefore, it is necessary to further assess the VOI predictor’s capability to predict VOI relationships across various insect species.

#### 3.2.1 Mixed dataset provides better features

Small-scale datasets often pose a risk of overfitting, as they fail to provide comprehensive feature information. To address this, we attempted to fuse existing experimental datasets to expand their scale and enhance the model’s predictive performance. As depicted in Fig. 2a, four datasets were combined using different strategies: Single, Ag-Dm, and Mix. Five-fold cross-validation was performed to evaluate model performance on each test set, with results for the single-dataset (Single) strategy previously reported (Fig. 1b). For the two mixed datasets (Ag-Dm and Mix), docking pretrain models showed more significant performance improvement than random init under all three partitioning scenarios (Fig. 2b).

**Fig. 2.**
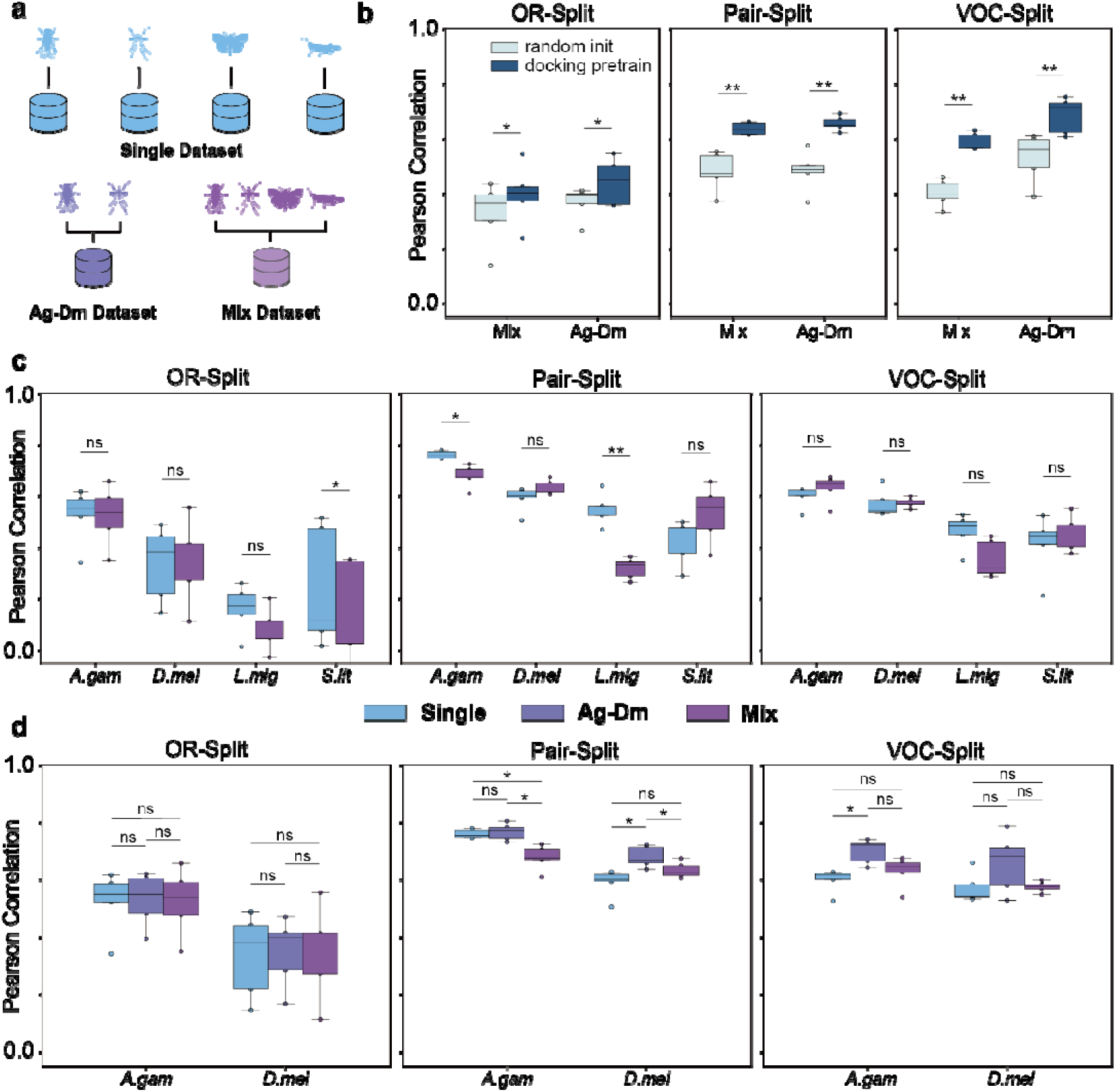
Performance of VOI predictor. a, Three data mixing schemes: Single (each species separately), Ag-Dm (same order combined), and Mix (all data pooled). **b**, Model performance under different data partitioning scenarios for the two mixed datasets. **c,d,** Performance of models trained on different datasets, evaluated separately on each species in cross-validation test sets.

#### 3.2.2 The applicable scope of the data fusion strategy

Evaluating model performance directly on the mixed dataset test set may not adequately reflect the model’s predictive ability for each individual species. Therefore, we split the test set data by species and calculated evaluation metrics separately for each species, with results shown in Fig. 2c-d. We found that for *Drosophila melanogaster* and *Anopheles gambiae*, the overall performance of the Ag[Dm model was better than that of the Single model (Fig. 2d; significant improvement was observed under Pair[Split and VOC[Split, and a higher median was achieved under the most stringent OR[Split), indicating that features from ORs of different species help improve the generalization performance of the model for cross[species VOI prediction. However, dataset fusion was not universally effective. As shown in Fig. 2c, the Mix model performed even worse than the Single model in most cases. We suppose that data from *Spodoptera littoralis* (Lepidoptera) and *Locusta migratoria* (Orthoptera) differ substantially from those of Diptera species (*Drosophila melanogaster* and *Anopheles gambiae*), limiting their ability to provide transferable features. Despite these variations, hybrid models consistently outperformed single-dataset models in species-specific predictions. Thus, we selected the Ag-Dm model as the final predictive framework, with its performance validated experimentally in subsequent sections.

### 3.3 Validation of the predictive capability of insectOlf

#### 3.3.1 Direct validation I: benchmarking against existing methods

We benchmarked insectOlf against several common machine learning methods and a sequence similarity-based approach(56). As shown in Tab. S2 and Fig. S7, insectOlf exhibited superior predictive performance on nearly all tested datasets.

#### 3.3.2 Direct validation II: functional experiments on an independent species

We further validated insectOlf’s cross-species predictive capacity through functional experiments on *Bactrocera dorsalis*, an independent pest species not included in the training data. We selected 6 ORs and 17 VOCs, yielding 102 OR-VOC pairs for testing; 10 positive interactions were identified, 2 of which have been previously reported(54) (concentration-response curves in Fig. 3c).

**Fig. 3.**
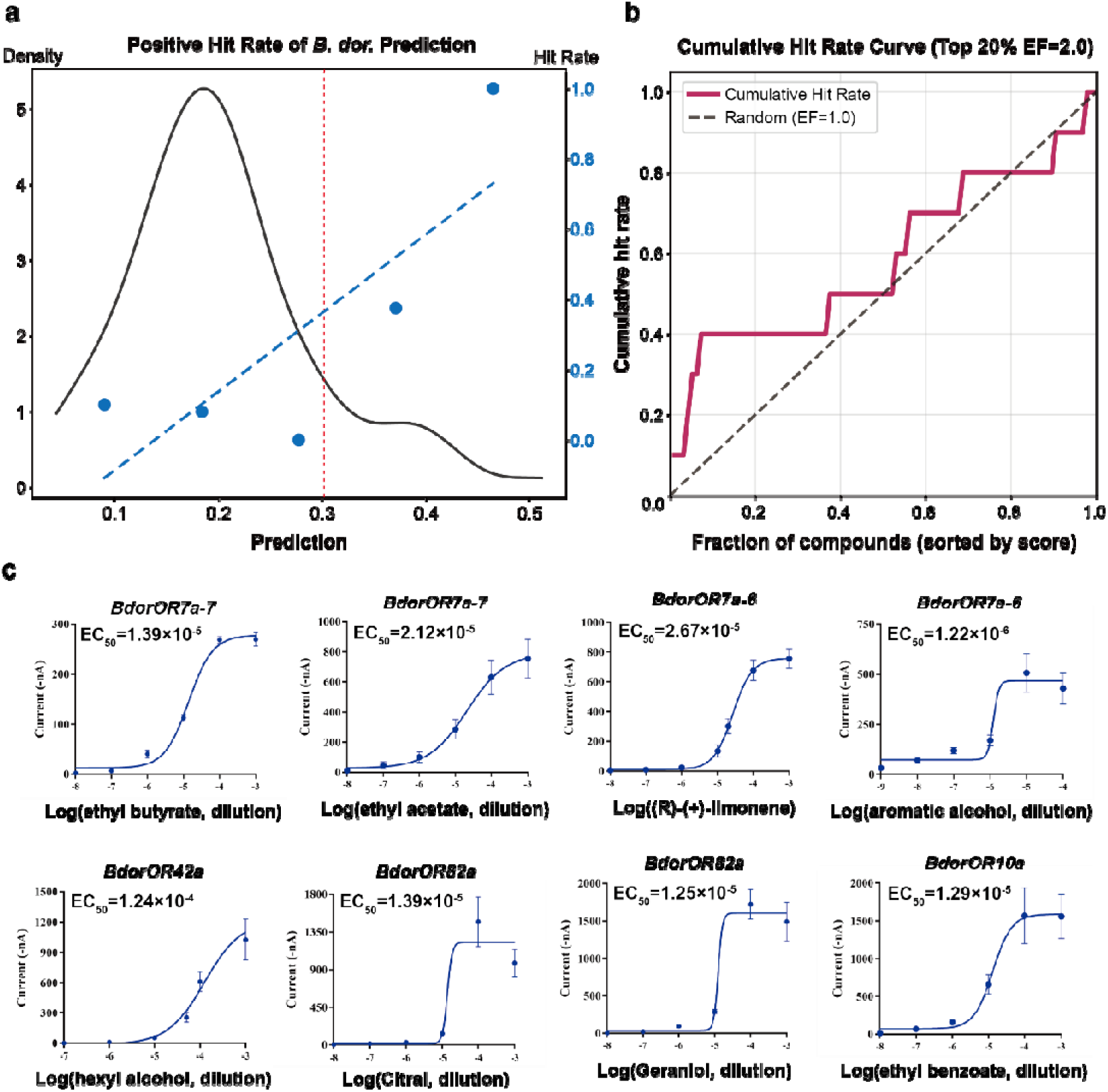
Validation of cross-species prediction via functional experiments on *Bactrocera dorsalis*. a, Prediction score distribution and hit rate of experimentally confirmed positives. **b,** Cumulative hit rate and enrichment factor of predictions. **c,** Concentration-response curves of the positive interactions.

Prediction scores for the 102 pairs are shown in Fig. 3a. When the score exceeded 0.3 (corresponding to the experimental threshold of spikes > 50), the hit rate of true positives increased markedly. Cumulative Hit Rate (CHR) and Enrichment Factor (EF) analyses (Fig. 3b) further confirmed that insectOlf enriches true positives among top-ranked predictions, with an EF of 2 in the top 20% of the data—twice the random expectation. We further tested insectOlf on a previously published larval *D. melanogaster* OR dataset(57) and observed enrichment consistently above random expectation (Fig. S17).

#### 3.3.3 Applicability and limitations

The cross-validation results on *D. melanogaster* and *A. gambiae*, together with the functional validation on *B. dorsalis*, collectively demonstrate insectOlf’s predictive capability within Diptera. To further explore the benefits of our transfer learning strategy, we simulated extreme data scarcity (20% training, 80% testing). The results confirmed that docking-based pretraining and data fusion from closely related species enhance predictive performance under such conditions (Fig. S8). However, cross-order predictions (e.g., to Lepidoptera or Orthoptera) performed poorly (Fig. S9), defining the current applicability boundary of insectOlf.

#### 3.3.4 Indirect validation I: recapitulation of known olfactory preferences

As an indirect validation, we asked whether insectOlf could recapitulate a well-established olfactory trend: *D. melanogaster* preferentially responds to esters, while *A. gambiae* shows stronger responses to aromatics(6). Using insectOlf, we predicted VOI scores for 11 fruit fly and 10 mosquito species against 14,499 VOCs, and calculated PBI scores for each species. As shown in Fig. 4a, the predictions faithfully reproduced the known preferences: fruit flies exhibited stronger ester recognition, whereas mosquitoes showed greater affinity for aromatics. PGLS analysis confirmed that this pattern remains highly significant after accounting for phylogenetic relatedness (p < 0.001; Fig. S14).

**Fig. 4.**
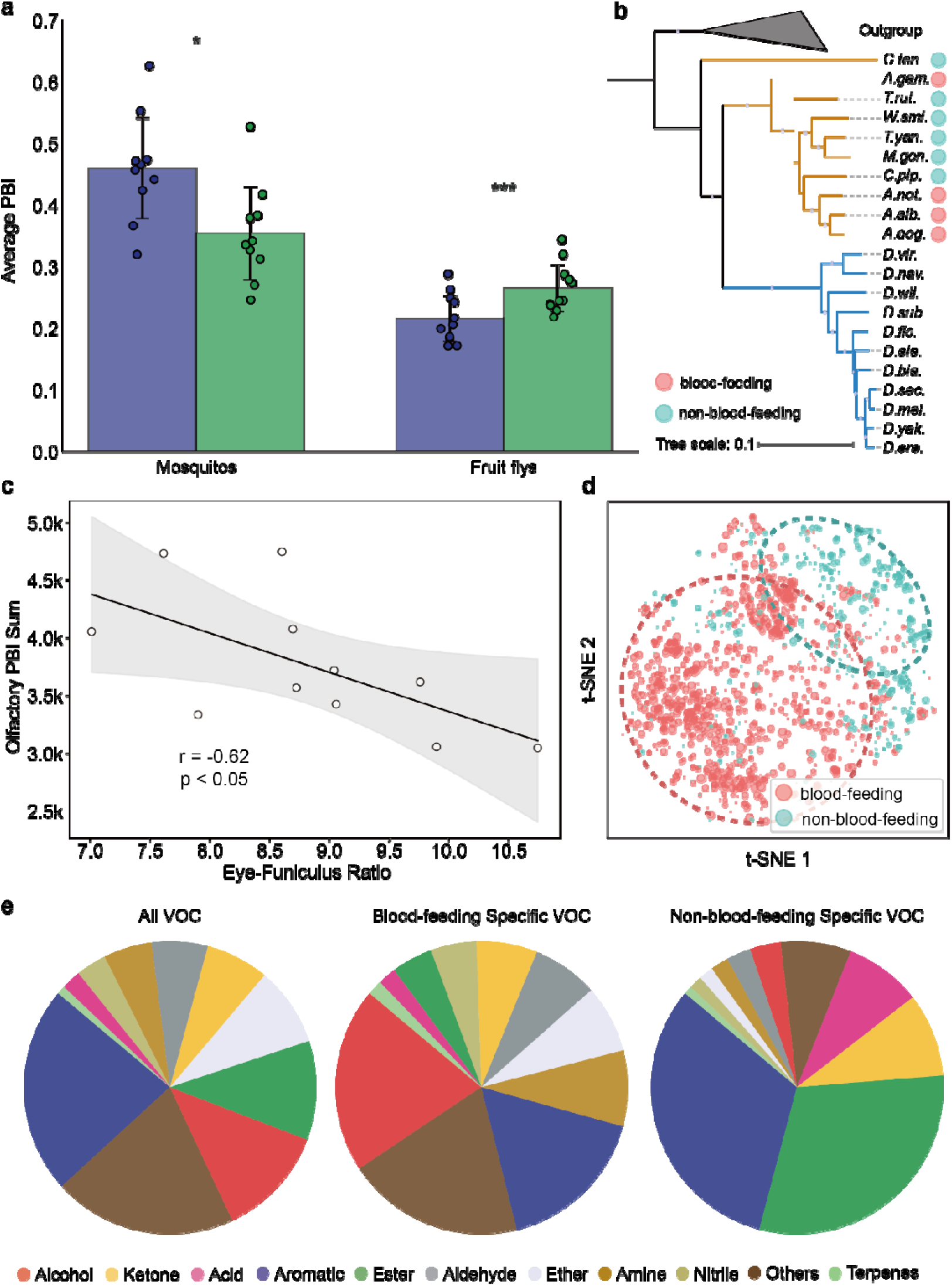
Application of insectOlf to olfactory preference analysis. a, PBI score differences between mosquitoes and fruit flies for aromatic and ester compounds. **b,** Phylogenetic relationships of species included in this study. **c,** Negative correlation between olfactory potential (PBI sum) and visual capacity (Eye-Funiculus Ratio) across Drosophila species. **d,** PBI score distribution of VOCs uniquely recognized by hematophagous vs. non-hematophagous mosquitoes. **e,** Functional group composition of VOCs across different datasets.

#### 3.3.5 Indirect validation II: recapitulation of the olfactory-visual trade-off

A second indirect validation tested whether insectOlf could reproduce the previously reported negative correlation between olfactory and visual capacities in Drosophila at the morphological level(58). We predicted VOI scores for the same 11 Drosophila species against 14,499 VOCs and calculated PBI-based olfactory potential. As shown in Fig. 4c, a significant negative correlation was observed between olfactory potential and visual capability (Eye-Funiculus Ratio), while OR count alone did not yield a significant correlation (Fig. S12). PGLS analysis confirmed this negative correlation remains significant after controlling for phylogeny (p < 0.05; Fig. S15).

### 3.4 Case study: olfactory specialization in mosquitoes with different feeding habits

Mosquitoes with different feeding habits (hematophagous vs. non-hematophagous) are expected to exhibit distinct olfactory sensitivities. Although the VOCs recognized by their ORs largely occupy similar chemical spaces (Fig. S16), clear differences emerged when we examined the unique VOCs specifically recognized by each group (Fig. 4d, bubble size represents PBI scores). Functional group analysis of these unique VOCs (Fig. 4e) revealed that, relative to the baseline distribution in the compound library, blood-feeding mosquitoes showed increased representation of alcohols, amines, and aldehydes—compounds often associated with host-derived odors—while aromatic compounds also remained prominent. In contrast, non-blood-feeding mosquitoes exhibited a marked increase in esters, consistent with plant-associated volatiles. These distinct chemical preferences suggest that host-seeking behavior in blood-feeding mosquitoes may rely on specific compound classes, providing a basis for prioritizing candidate molecules in vector behavioral studies.

## 4 Discussion

Molecular docking provides a scalable means to generate affinity scores between proteins and compounds, though its accuracy is constrained by structure prediction reliability and algorithmic limitations(59, 60). Nevertheless, docking outputs offer stable, moderately precise features across broad chemical spaces without relying on specific training datasets(25). In VOI prediction, high-dimensional features paired with sparse experimental data risk overfitting, whereas large-scale docking data provide a global view of the feature space that mitigates this risk. Our transfer learning strategy, which integrates docking-derived global features with experimental measurements, offers a viable pathway for enhancing model generalization.

We further evaluated the performance boundaries of insectOlf. The model achieved reliable cross-species prediction within Diptera (e.g., between *D. melanogaster* and *A. gambiae*), but performance declined markedly for cross-order predictions (Fig. S9). Data fusion from distantly related species did not consistently improve performance, suggesting that while certain functional patterns may be conserved across species(61), substantial divergence limits transferability. These findings indicate that insectOlf’s generalization capability correlates with phylogenetic distance, defining its current scope of application within Diptera.

Within this applicable scope, insectOlf provides a validated computational pipeline for accelerating the discovery of behaviorally active semiochemicals. The successful experimental validation of its top[ranked predictions for *B. dorsalis* (Fig. 3) demonstrates its potential to dramatically reduce the number of compounds requiring labor[intensive behavioral screening by generating a prioritized shortlist. For a pest species of interest, researchers could use insectOlf to screen thousands of semiochemicals, obtain a ranked list of predicted activators for its OR repertoire, and then focus behavioral assays (e.g., olfactometer, field trapping) on the top candidates, thereby streamlining the discovery pipeline for novel attractants or repellents.

Beyond direct prediction, insectOlf’s ability to recapitulate established ecological patterns further supports its utility for informing pest management strategies. The framework accurately captured the distinct ester/aromatic preferences between Drosophila and mosquitoes (Fig. 4a), confirming that it can identify chemical vulnerabilities characteristic of a pest lineage—information that can guide the selection of candidate semiochemicals for broad[spectrum monitoring traps. Moreover, the recovery of the sensory trade[off between olfactory and visual investment in Drosophila (Fig. 4c) illustrates how insectOlf can infer a species’ relative reliance on olfaction, a key factor in assessing the likely efficacy of odor[based control tactics. By translating genomic and chemical data into these ecologically meaningful insights, insectOlf offers a computational foundation for rational design of species[or lineage[targeted attractants and for prioritizing pest systems where olfactory intervention holds greatest promise.

As a proof-of-concept application, we used insectOlf to explore olfactory specialization in mosquitoes with different feeding habits. Although the chemical spaces recognized by hematophagous and non-hematophagous mosquitoes largely overlap, clear differences emerged in the VOCs uniquely recognized by each group. Blood-feeding mosquitoes showed increased representation of alcohols, amines, and aldehydes—compounds often associated with host-derived cues—while non-hematophagous mosquitoes exhibited a marked enrichment of esters, consistent with plant-associated volatiles. This pattern suggests that host-seeking behavior in blood-feeding mosquitoes may rely on specific chemical classes, offering a basis for prioritizing candidate molecules in vector control research. More generally, our framework enables the systematic identification of chemical features associated with distinct ecological traits, which could guide the development of species-specific attractants or repellents by narrowing the chemical space that warrants further behavioral testing.

One methodological caveat concerns the PBI parameter used to quantify olfactory potential. PBI is suitable for assessing broad preferences (e.g., towards host-associated vs. plant-associated volatiles) because these recognition processes often involve many[to[many VOI relationships(62–64). However, PBI is less informative for narrow[spectrum interactions such as pheromone detection, where strong responses to specific molecules may yield low PBI values. This limitation should be considered when interpreting our results and highlights an area for future methodological refinement.

In summary, we have developed insectOlf, a transfer learning-based framework that integrates large-scale molecular docking with sparse experimental data to predict OR-VOC interactions. The framework was validated through both cross-validation and functional experiments on an independent pest species, and its utility was demonstrated through a case study identifying distinct chemical preference patterns between hematophagous and non-hematophagous mosquitoes. Looking forward, by enabling systematic comparison of olfactory preference patterns across species, insectOlf can inform the prioritization of chemical libraries for behavioral screening and may assist in identifying ORs associated with host-seeking or other behaviors, thereby facilitating target selection for subsequent functional studies. Ultimately, approaches like this could accelerate the discovery of behaviorally active compounds, contributing to the development of environmentally sustainable pest management strategies. That said, the current model is most reliable within Diptera, and cross-order predictions remain challenging. Expanding the coverage of experimental OR functional data across diverse taxa will be essential for improving generalization, much as comprehensive structural databases underpin advances in protein structure prediction. We hope insectOlf serves as a useful computational resource for researchers working on insect olfaction and behavioral control. By enabling the rapid, in silico profiling of pest olfactory landscapes, insectOlf has the potential to lower the barrier to entry for semiochemical discovery, empower the design of precision attract-and-kill systems, and contribute to the development of more sustainable and effective pest management protocols.

## Ethics approval

The animal experiments were conducted under the license of the Animal Care and Use Committee of the Southwest university (IACUC-20221114-04). *Xenopus laevis* was anesthetized 30 min by bathed in the ice, and wounds were carefully treated to minimize suffering and avoid infection.

## Code availability

**Data, model, and scripts of insectOlf download address:** GitHub (https://github.com/iORbase/insectOlf).

## Acknowledgements

This work was supported, in part, by the National Natural Science Foundation of China (32270525, 32470445); the Pherobio Semiochemical Institute Open Program (PHROBIO2023ZJSF02); the Natural Science Basic Research Program of Shaanxi (2021JM-212). We are grateful to Professor Chenzhu Wang from Shanxi University for his valuable guidance and constructive comments during the revision of this manuscript.

## Author contributions

Y.Z. and L.X. contributed equally. The research was designed by H.L. and H.J. and polished by C.X and G.L. The literature data was collected and analyzed by T.Z., C.X., Q.S., and G.L. The silico experiments were performed and analyzed by Y.Z., C.G., S.D., Y.Y., X.Q. and X.Y. The in vitro experiments were performed and analyzed by L.X. The paper was written by Y.Z. and H.L. All authors have given approval to the final version of the manuscript.

## Supporting information

The online version contains supplementary material available. Correspondence and requests for materials should be addressed to Huimeng Lu.

## Data availability

The supplementary materials contain data that support this study’s findings, and more information is available from the corresponding authors upon request.

