## supplement text for "Transfer learning reveals species-specific olfactory preferences in Diptera and informs pest management strategies"

**Insect OR Gene Annotation**

Insect genomes for olfactory potential and environmental adaptation analysis were downloaded from the NCBI public database (including 19 *Drosophila* species, 4 blood-feeding mosquitoes, and 7 non-blood-feeding mosquitoes). We employed insectOR^1^ to annotate OR genes from these genomes, followed by rapid structure prediction using the iModelTM module of iORbase^2^. Structural screening was performed using Dali^3^, retaining only structurally intact, functional ORs. Ultimately, we retained OR repertoires from 11 *Drosophila* species with reasonable OR counts and available visual information, along with those from 4 blood-feeding and 6 non-blood-feeding mosquitoes. Detailed genome information is provided in the supplementary materials.

**Functional group classification of compounds**

For the experimental data used to train the model, the functional group information of VOCs was derived from the labels in the original publications. For the large-scale compound dataset we used, functional group labels were computed using RDKit, and the final functional group label for each molecule was determined according to the functional group priority rules specified in IUPAC standards^4^ (e. g. :carboxylic acid > ester > aldehyde > ketone > alcohol > amine > aromatic > hydrocarbon > others).

### A predictive framework for insect olfactory potential in environmental adaptability analysis

*InsectOlf workflow.* As illustrated in Fig. S1, we constructed a predictive framework (insectOlf) for evaluating insect olfactory ability and environmental adaptability. Specifically, based on the transfer learning strategy mentioned above, we developed a VOI predictor capable of batch predicting the VOI relationships between the OR repertoire of target insects and a large library of VOCs. Subsequently, we calculated the potential binding index (PBI) of target insects using these VOI relationships to assess the olfactory ability of target insects. The PBI has previously been applied in functional evolution and environmental adaptability analyses^5^.

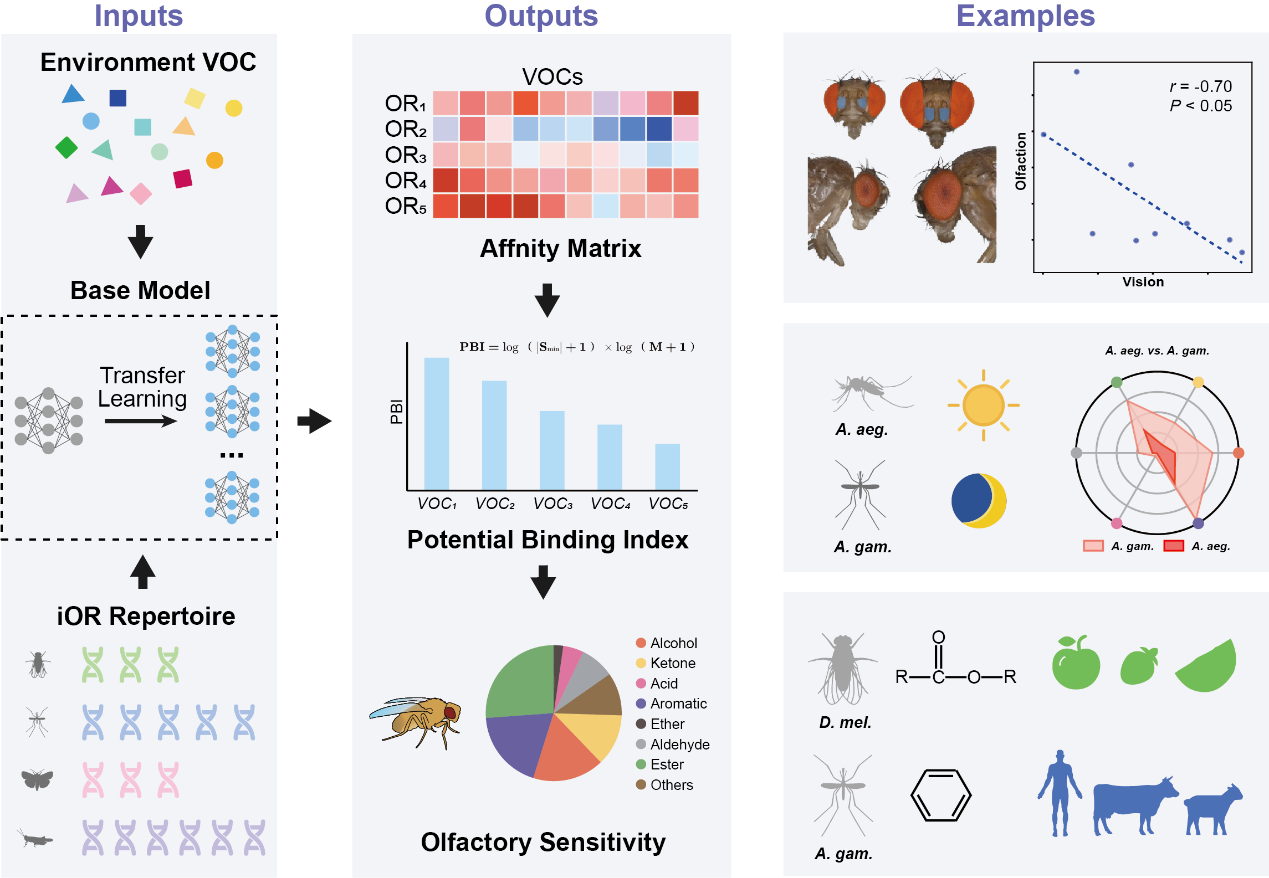

**Fig. S1 | insectOlf workflow**

### Model architecture for VOI prediction

Protein sequences were encoded using the ProtT5-XL-UniRef50 model(https://huggingface.co/Rostlab/prot_t5_xl_uniref50), and small-molecule SMILES expressions were encoded via a pre-trained Roberta model(https://huggingface.co/DeepChem/SmilesTokenizer_PubChem_1M). The feature vectors from both sources were concatenated into a 1792-dimensional feature vector, which was then fed into a fully connected neural network to output the corresponding prediction scores. Relevant parameters of the model are provided in Table S1, and the architecture is illustrated in Fig. S2.

**Table S1: the hyper-parameters of neural network model**

| Hyper-parameters | Value |
| --- | --- |
| number of nodes for input layer | 1792 |
| number of nodes for fully connected layer | 512 |
| number of nodes for batch normalization layer | 512 |
| Optimizer | Adam |
| initial learning rate | 10^-4^ |
| Learning rate scheduler | ReduceLROnPlateau  (factor=0.5, patience=5) |
| max epoch | 50 |
| min epoch | 20 |
| early stopping patience | 10 |
| batch size | 512 |
| seed for train_test_split | 1234 |

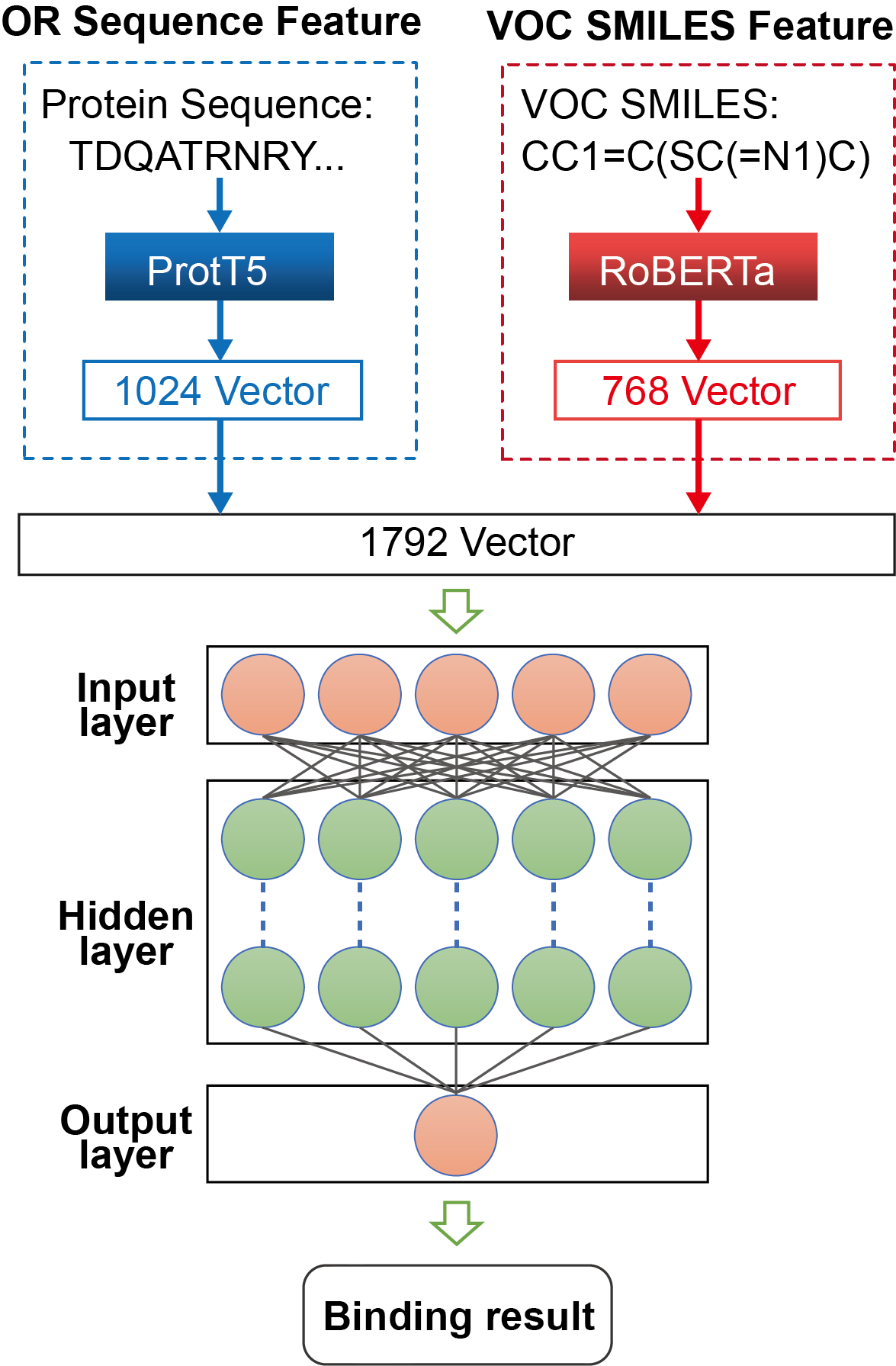

**Fig. S2 | Model architecture for VOI prediction**

### Docking data expands the chemical space

The docking dataset we used contains 2,077 pheromones closely associated with various physiological activities of insects. Its scale far exceeds the number of VOCs used in any existing experimental dataset, as illustrated in Fig. S3. More VOCs provide more comprehensive feature representations and cover a broader chemical space, as shown in Fig. S4.

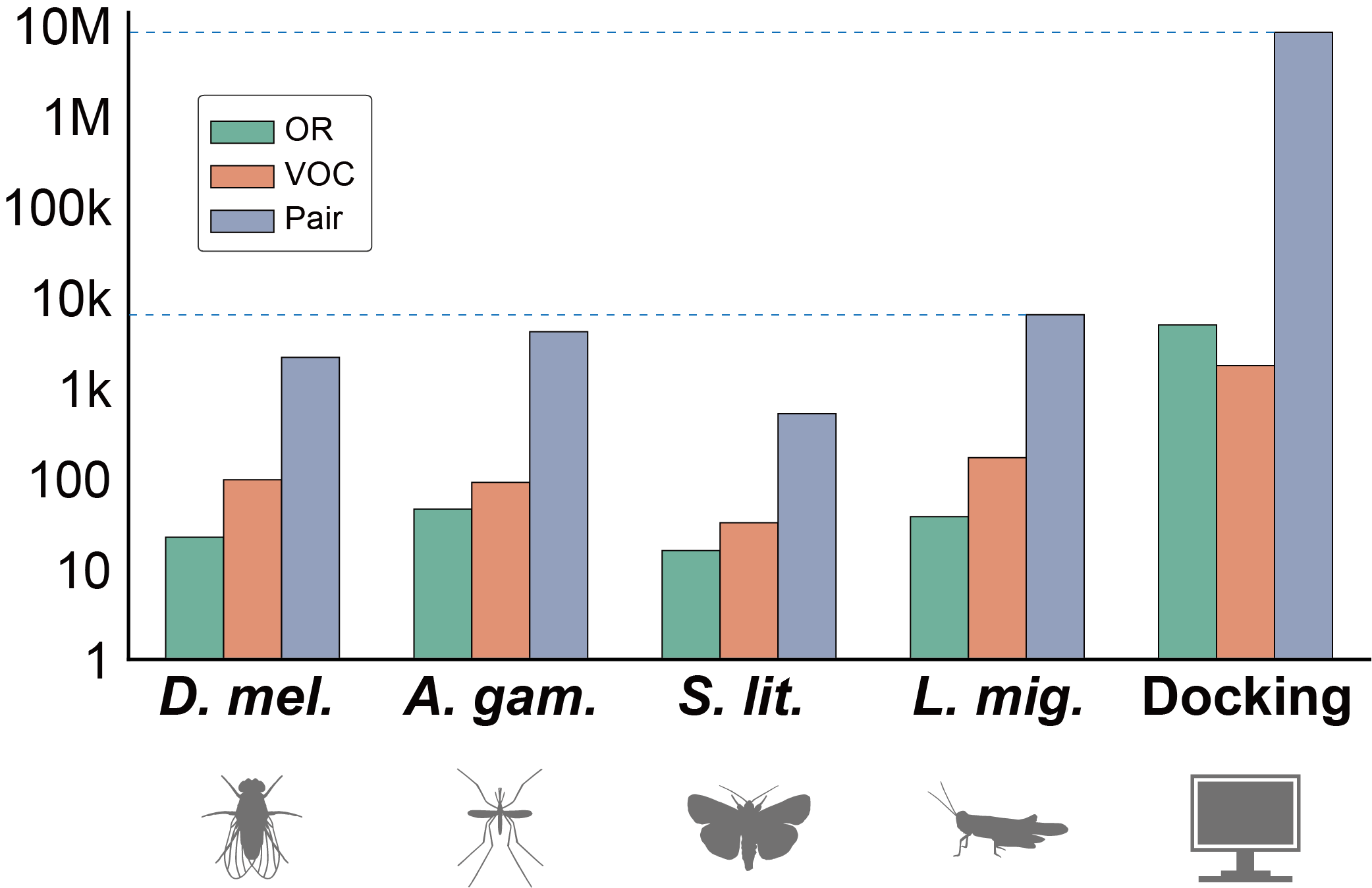

**Fig. S3 | Statistics on the quantity of different datasets**

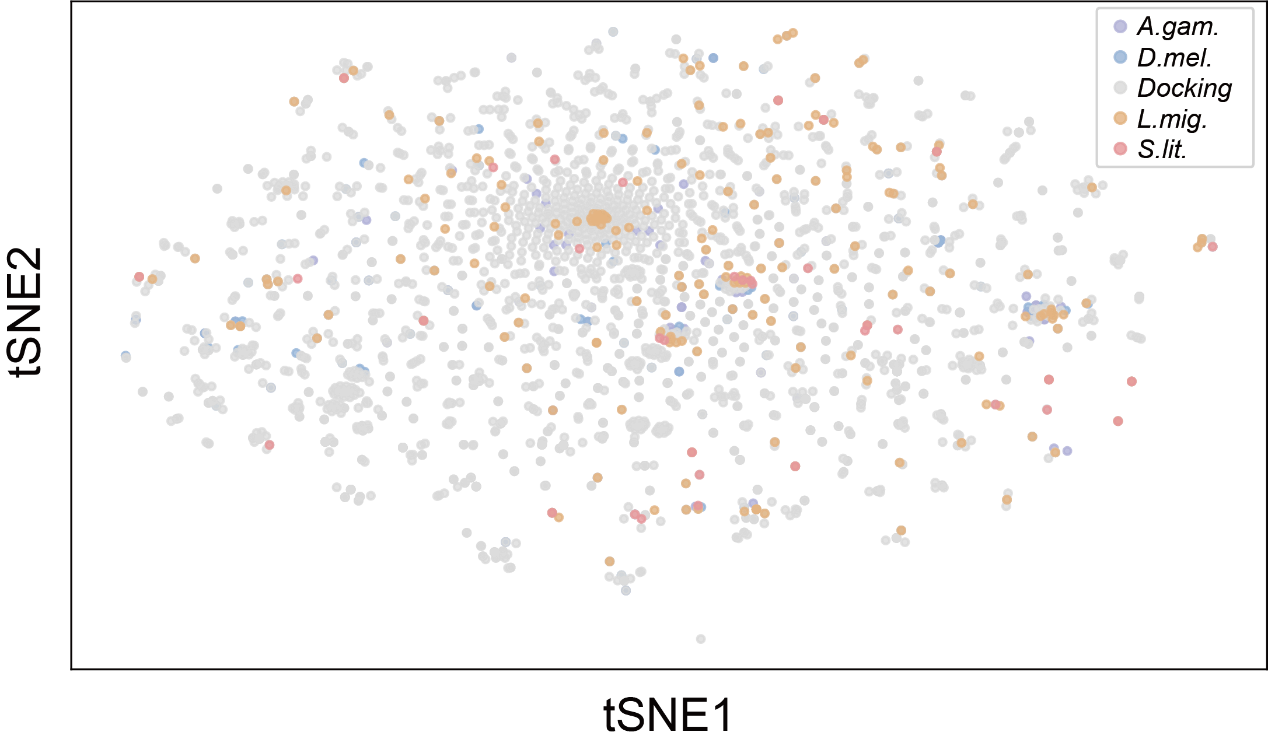

**Fig. S4 | Feature Distribution of Different Datasets**

### Consistency between molecular docking methods and real experimental data

Taking the fruit fly dataset as an example, we evaluated the screening performance of molecular docking methods (exemplified by Vina) for positive results in real experiments. As shown in Figure S5, the blue curve represents the distribution of docking scores, while the red points indicate the hit rate of positive results within each bin after equally spaced binning of all docking scores (i.e., the proportion of true positive results in the data points of that bin). It can be observed that as the absolute value of the docking score increases (docking scores greater than zero are considered meaningless), the hit rate of experimentally validated positive results gradually increases, reaching its highest value in the bin with the highest docking scores (approximately 30%, which is generally consistent with hit rates reported in previous large-scale compound screening workflows^6^). This indicates that molecular docking methods can enrich true positive results to a certain extent and provide an overall VOI distribution trend for transfer learning. However, due to the presence of a large number of false positives, conventional classification evaluation metrics are not suitable for assessing the performance of molecular docking methods. Therefore, we did not compare them directly with our model.

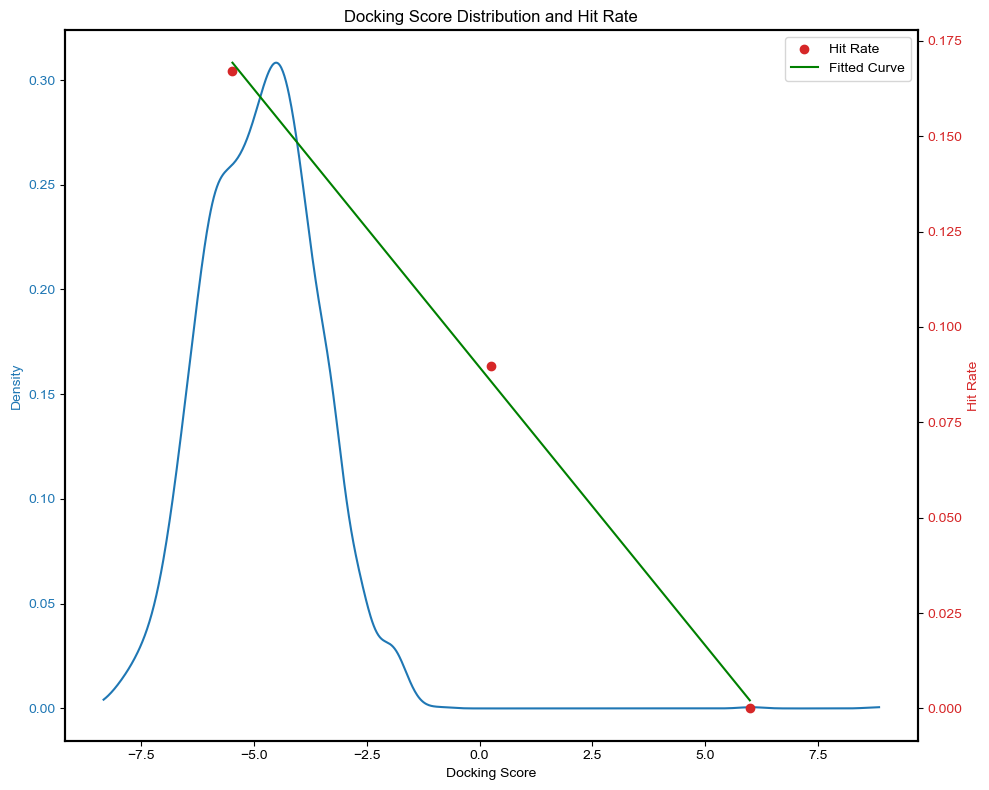

**Fig. S5 | The hit rate distribution of the Vina method on the *D. mel.* dataset**

### Consistency of different docking methods

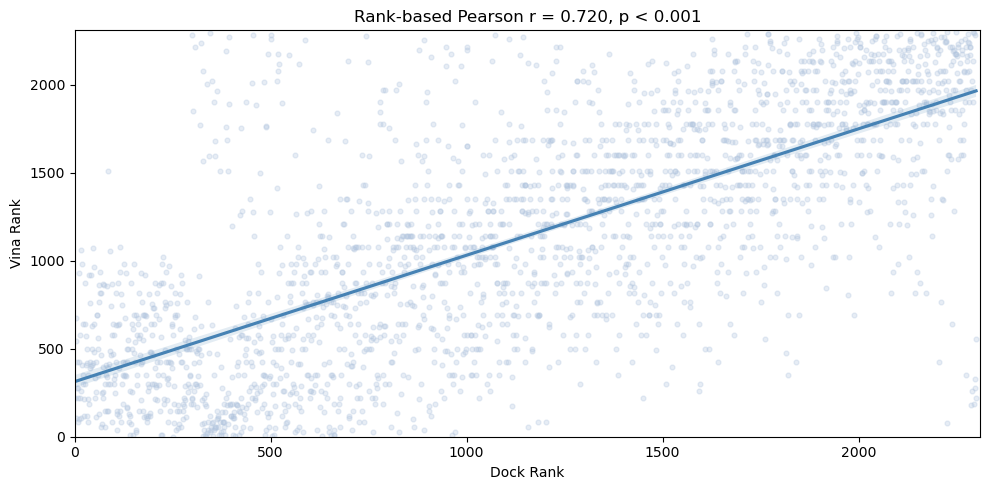

**Fig. S6 | Consistency of Vina and DOCK results**

To assess whether there are significant differences between different docking methods, we evaluated the consistency between the docking results of Vina and DOCK (both methods were applied to OR-VOC docking in the *Drosophila melanogaster* and *Anopheles gambiae* experimental datasets; data that failed format conversion during preprocessing were excluded). The results are shown in Figure S6. There was a significant correlation between the rank results of the two molecular docking methods (Pearson r = 0.72, p < 0.001). Therefore, we consider that different docking methods do not have a disruptive impact on our transfer learning strategy.

### Comparison of VOI prediction methods

Using 4 datasets, we implemented several common machine learning algorithms for regression tasks and quantified the correlation coefficients between predicted and observed values. Results demonstrated that our model achieved superior predictive performance compared to benchmark methods, as shown in Tab. S2.

Chepurwar *et al.* proposed a method based on OR sequence similarity to predict VOIs of other ORs using existing data. We performed predictions on the same dataset using VOI predictor and compared the prediction performance against the results reported in their original study (Fig. S7).

**Table S2: Comparison of the performance of the regression task of the VOI prediction method**

| Model | R (↑) | | | | MSE (↓) | | | |
| --- | --- | --- | --- | --- | --- | --- | --- | --- |
|  | *D. mel* | *A. gam* | *L. mig* | *S. lit* | *D. mel* | *A. gam* | *L. mig* | *S. lit* |
| **VOI predictor** | **0.679(±0.037)** | **0.767(±0.030)** | **0.551(±0.062)** | **0.416(±0.085)** | **0.020(±0.002)** | **0.008(±0.001)** | **0.001(±0.000)** | **0.013(±0.004)** |
| KNN | 0.300(±0.018) | 0.485(±0.018) | 0.092(±0.021) | 0.274(±0.033) | 0.023(±0.003) | 0.011(±0.001) | 0.001(±0.000) | 0.012(±0.003) |
| LR | 0.563(±0.037) | 0.554(±0.020) | 0.510(±0.093) | 0.342(±0.080) | 0.016(±0.003) | 0.010(±0.001) | 0.001(±0.000) | 0.011(±0.002) |
| SVM | 0.491(±0.050) | 0.474(±0.015) | 0.306(±0.063) | 0.286(±0.073) | 0.019(±0.003) | 0.012(±0.002) | 0.008(±0.000) | 0.013(±0.002) |

**Table S3: Hyperparameters used in other methods**

| Model | Hyperparameter | Value |
| --- | --- | --- |
| KNN | n_neighbors | 10 |
|  | weights | uniform |
|  | algorithm | auto |
|  | metric | minkowski |
| LR | fit_intercept | True |
|  | copy_X | True |
| SVM | C | 1.0 |
|  | epsilon | 0.1 |
|  | kernel | rbf |
|  | gamma | scale |
|  | max_iter | -1 |

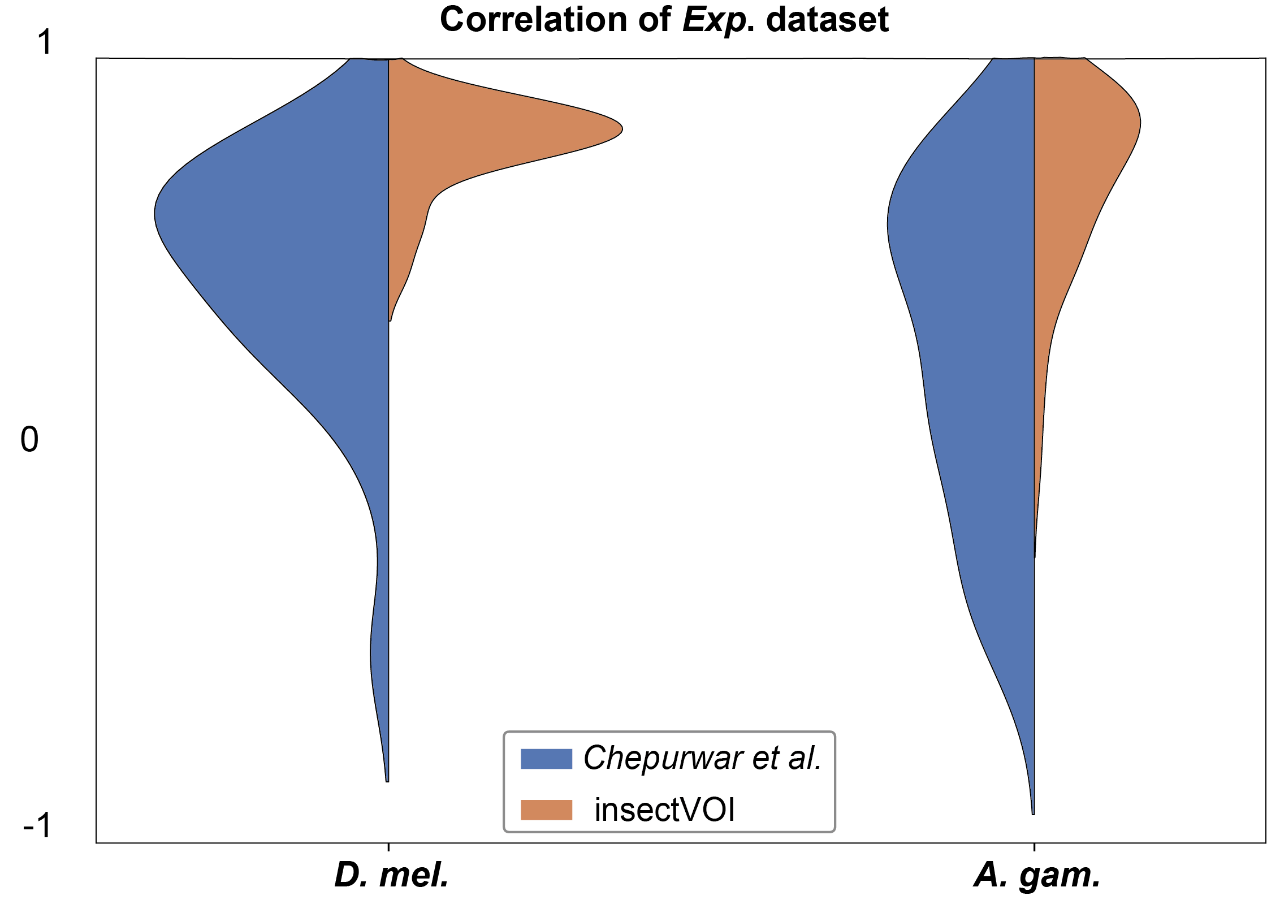

**Fig. S7 | The model performance of sequence-based method and VOI predictor**

### Performance under extremely scarce data conditions

For situations in which available data are insufficient, we compared 4 different methods: (1) Cross-species predict, in which the OR function of target species is directly predicted by the neighbor species-based model; (2) Directly training, train the model directly using the small amount of data that is already available; (3) Single-transfer, in which a tiny dataset of target species is used to refine the docking-based model and (4) Multi-transfer, in which the neighbor species-based model is refined.

For species with sparse experimental results, using existing data alone may not be sufficient to construct a functional prediction model. Here, we simulated the lack of available data on the *Drosophila melanogaster* datasets and compared the predictive capabilities of four different modes (Cross-predict, Directly train, Single-transfer, and Multi-transfer). The results indicate that transfer learning can significantly improve model performance, with the multi-transfer mode providing the best predictions. This demonstrates that the multi-transfer learning approach can effectively address the issue of insufficient existing data.

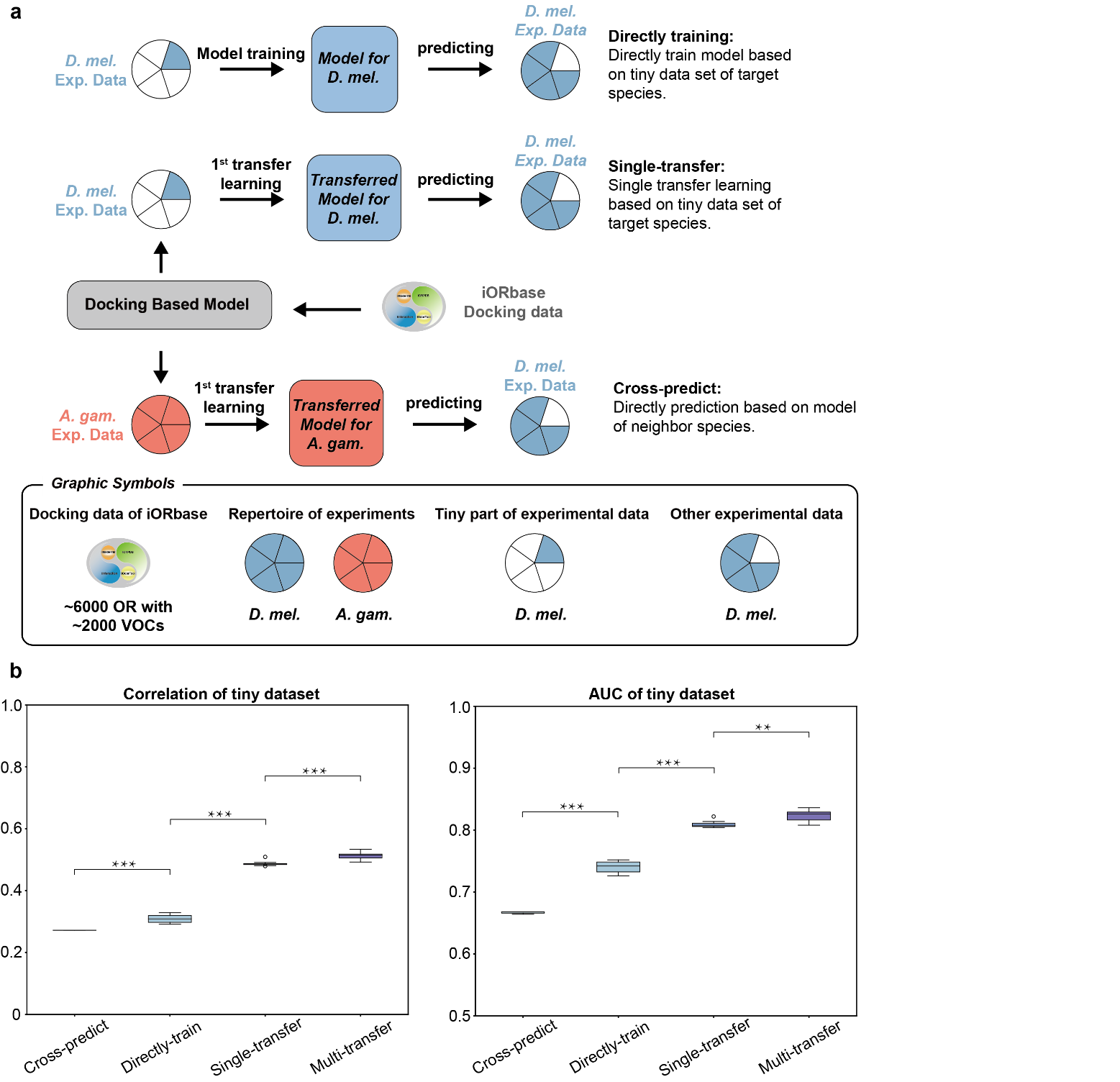

**Fig. S8 | Interspecific prediction performance analysis. a,** Using the fruit fly dataset as an example, 20% of the total data is designated as training data (tiny dataset) to mimic situations with limited available data. The figure illustrates the flow of OR function prediction for target species using four different modes. **b,** Compare the performance (correlation between experimental data and predicted values, AUC of prediction models) of four different modes (Cross-predict, Directly-train, Single-transfer and Multi-transfer) on the *D. melanogaster* tiny datasets. Results from the four modes significantly differ on the tiny datasets based on paired sample t-tests (** P < 0.01, *** *P* < 0.001).

### Cross-order predictions are often unreliable

To assess cross-species predictive ability, we used a model trained on *Drosophila melanogaster* and *Anopheles gambiae* (Ag-Dm model) to perform regression predictions of binding scores for two insect species not included in the training set: *Locusta migratoria* (*L. mig.*) and *Spodoptera litura* (*S. lit*)*.*

As can be clearly observed from the Fig. S9, regardless of how the true values change, the model outputs are highly concentrated in the 0.5–0.6 range. This phenomenon indicates that the model fails to identify the binding characteristics of the target species, unable to make effective distinctions, resulting in prediction results tending toward intermediate values.

Especially in the *L. mig.* data, despite the large sample size, the model predictions still failed to show effective correlation with the actual values, indicating that the model is almost unable to generalize to pair prediction tasks across different species. Therefore, direct transfer learning across species is unreliable in this task, emphasizing the significant impact of species differences on OR-VOC prediction models.

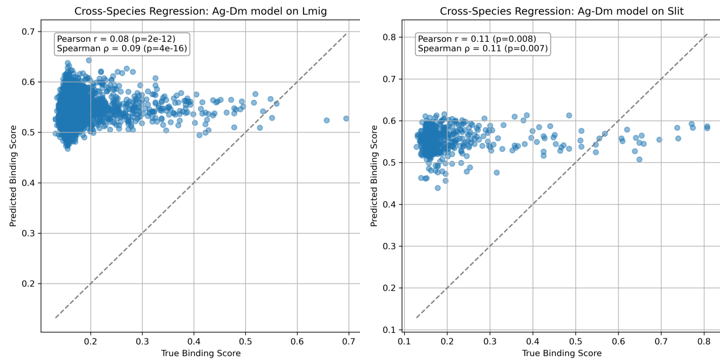

**Fig. S9 | The Predictions of Ag-Dm model on *L.mig.* and *S. lit.***

### Statistical results after removing experiment molecules

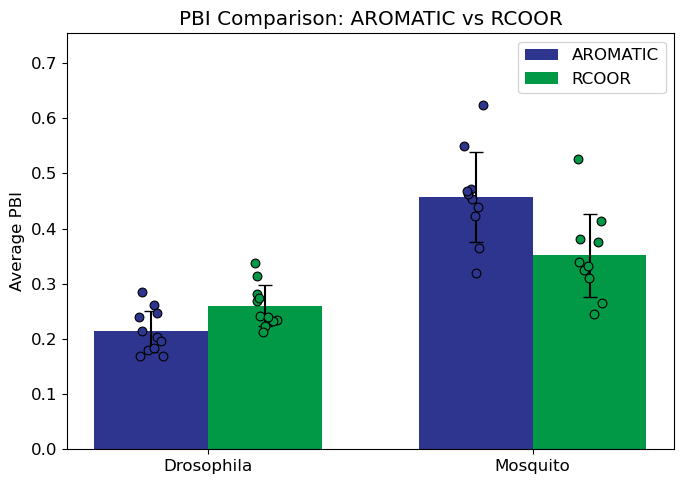

**Fig. S10 | Statistical results after removing experiment molecules**

To avoid potential misleading data bias, we removed from the 14,499 VOC dataset all VOCs (250 VOCs in total) that appeared in the experimental dataset used for model training, and re-calculated the preferences of fruit flies and mosquitoes for esters and aromatics, obtaining conclusions consistent with those previously drawn.

### The number of ORs between fruit flies and mosquitoes

We analyzed 11 fruit flies and 10 mosquitos from all annotation results, and found that the number of ORs in mosquitoes was significantly higher than that in fruit flies.

Additionally, the number of ORs in mosquitoes exhibited greater variation, which may be associated with differences in feeding habits (hematophagous vs. non-hematophagous) or their phylogenetic relationships.

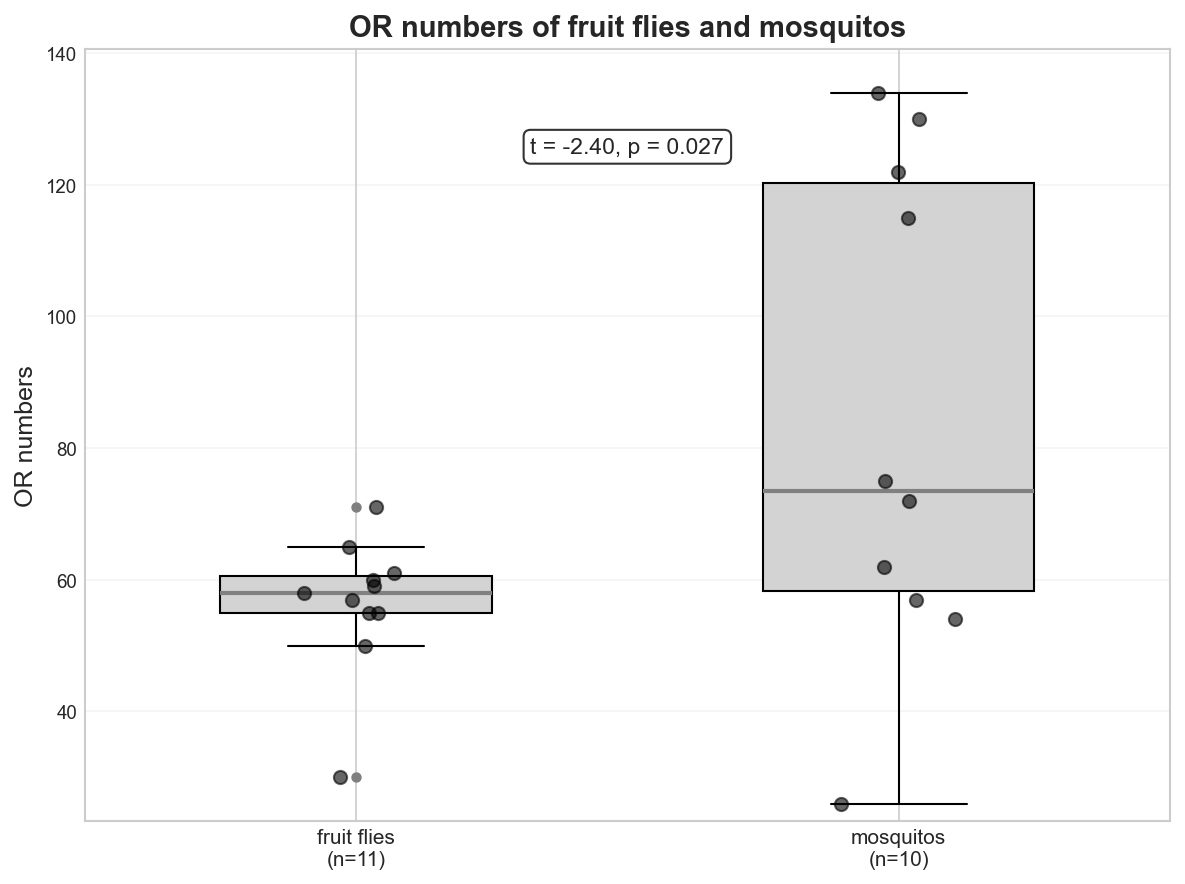

**Fig. S11 | The number of OR between fruit flies and mosquitoes**

### Negative correlation between vision and olfaction

We used the number of OR and the docking result of OR in Drosophila to analyze the correlation between the olfaction and the vision, respectively. We then found that there was no significant correlation between the number of OR or the docking results and visual ability in *Drosophila*.

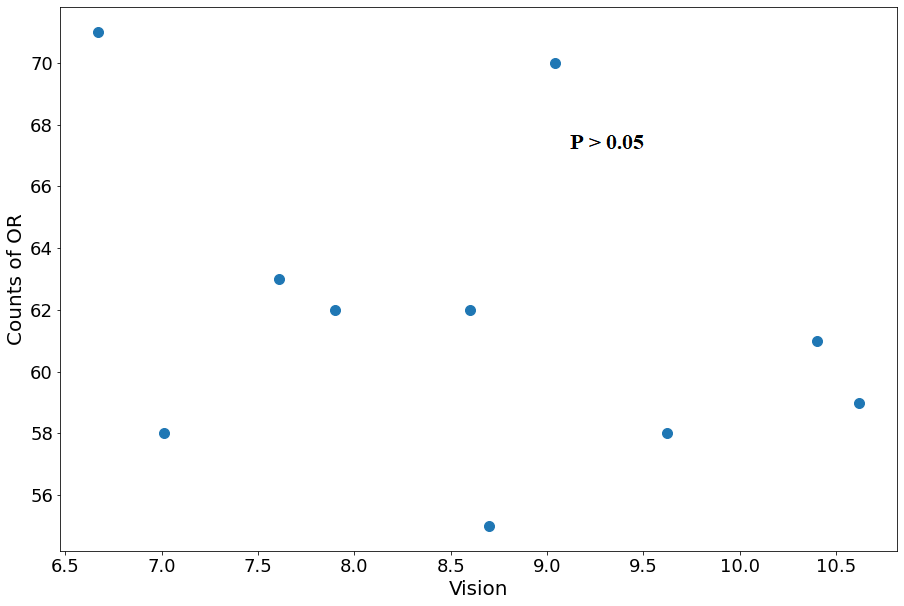

**Fig. S12 | The number of OR and visual ability in *Drosophila*.**

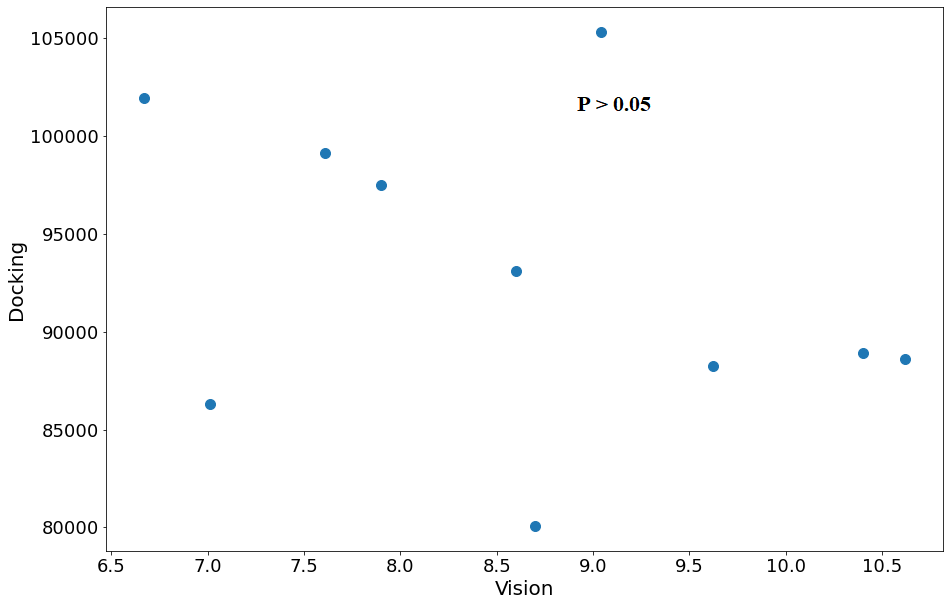

**Fig. S13 | Docking results and visual ability in *Drosophila*.**

### Phylogenetic tree construction and Phylogenetic Generalised Least Squares (PGLS) analysis

A total of 21 Diptera species were included in this study. In addition, four outgroup species (*Panorpa liui*, *Xenopsylla cheopis*, *Ctenocephalides felis*, and *Panorpa germanica*) were selected for rooting the phylogenetic tree. Genomic data for all species were obtained from public databases (NCBI). The calculation methods for trait data (including ester_PBI, aromatic_PBI, EF-ratio, and PBI_sum) have been described in the main text.

BUSCO (v5.8.2) was used with the metaeuk mode and the dipter_odb12 database (2025-07-01 release) to identify single-copy orthologous genes in each genome. Using busco2fasta.py (available at https://github.com/lstevens17/busco2fasta), we identified 3,095 single-copy BUSCO genes present in at least 90% of the genomes. Protein sequences of these BUSCOs were aligned using MAFFT (v7.525), and trimAl (v1.4) was used to trim the alignments, retaining conserved regions and removing ambiguously aligned sites. A total of 2,596 alignments passed the quality threshold. The trimmed alignments were concatenated into a supermatrix using catfasta2phyml (available at https://github.com/nylander/catfasta2phyml). This supermatrix was provided to IQ-TREE (v2.03) to infer the species tree under the LG+G model, with 1,000 ultrafast bootstrap replicates^7^. Finally, the tree was rooted at the common ancestor node of Diptera and the outgroup species using iTOL v6^8^, obtaining a rooted tree for subsequent analyses.

Given that trait data across species are non-independent in statistical analyses due to shared evolutionary history, direct use of conventional regression may lead to spurious associations. Phylogenetic Generalised Least Squares accounts for this non-independence by incorporating a phylogenetic covariance matrix, thereby providing more accurate estimates of evolutionary correlations between variables. The PGLS model was implemented using the caper package in the R environment^9^.

### Results of PGLS analysis

To compare olfactory preferences between fruit flies and mosquitoes while controlling for phylogenetic relationships, a PGLS model was fitted with species group (Drosophila vs. Culicidae) as the independent variable and the difference between ester PBI and aromatic PBI as the dependent variable. The analysis revealed a highly significant difference between the two groups (p < 0.001), indicating that fruit flies’ relative preference for esters and mosquitoes’ relative preference for aromatics are not simply inherited from a common ancestor but represent adaptive traits linked to their respective ecological niches.

Similarly, a PGLS regression was performed across 11 Drosophila species to test the negative correlation between visual and olfactory capabilities, using PBI_sum as the dependent variable and EF-ratio as the independent variable. After accounting for phylogenetic relationships, the negative correlation remained significant (regression coefficient t-test p < 0.05), confirming that this trade-off is not an artifact of shared ancestry.

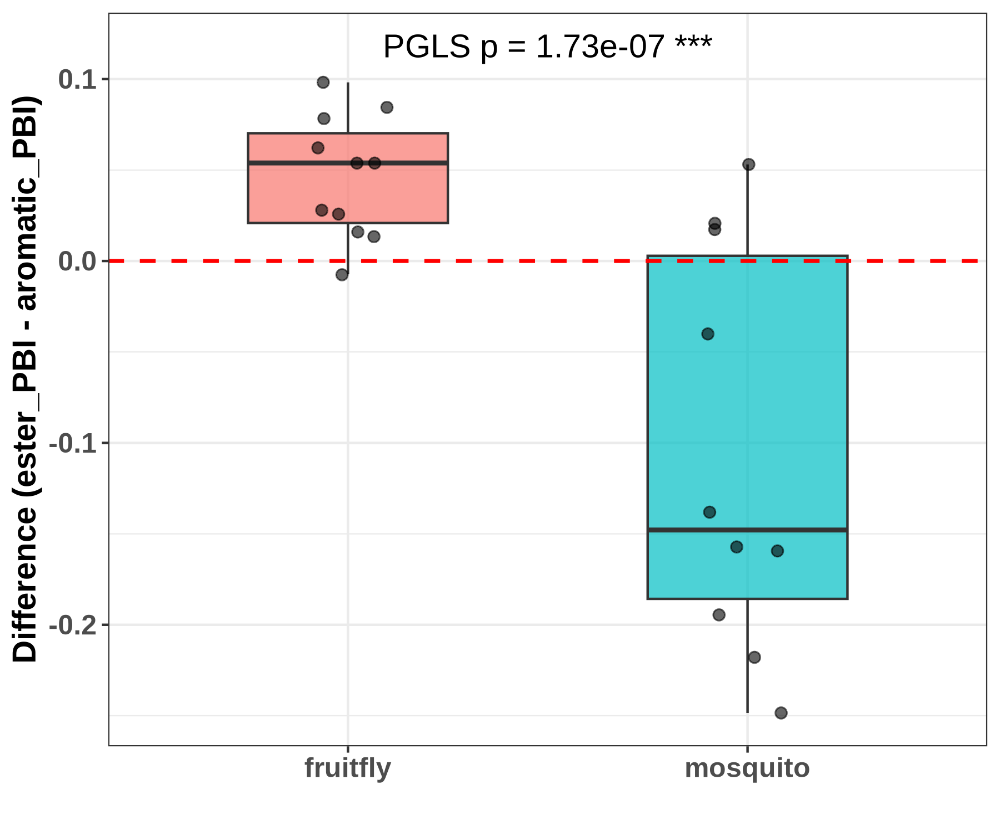

**Fig. S14 | PGLS result of Olfactory divergence between fruit flies and mosquitoes**

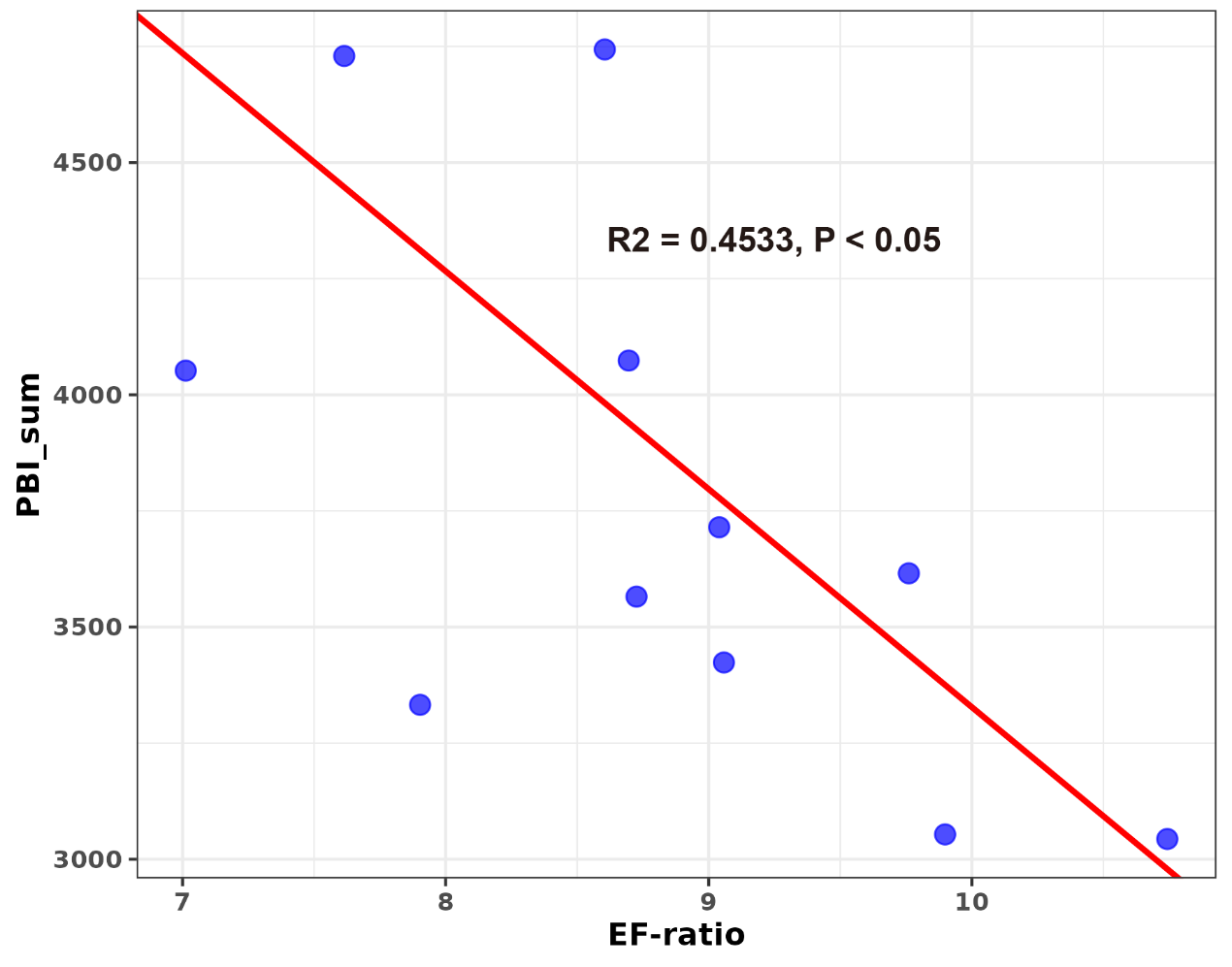

**Fig. S15 | PGLS result of Olfactory-visual correlation in *Drosophila*.**

### The chemical space recognized by hematophagous and non-hematophagous mosquitoes is similar

As shown in Fig. S16, the VOCs recognized by the ORs of blood-feeding and non-blood-feeding mosquitoes largely occupy similar chemical spaces. However, the distribution of the proportion of strong-binding VOCs (indicated by bubble size, representing the fraction of ORs with strong responses among all ORs responding to VOCs) shows discernible variations.

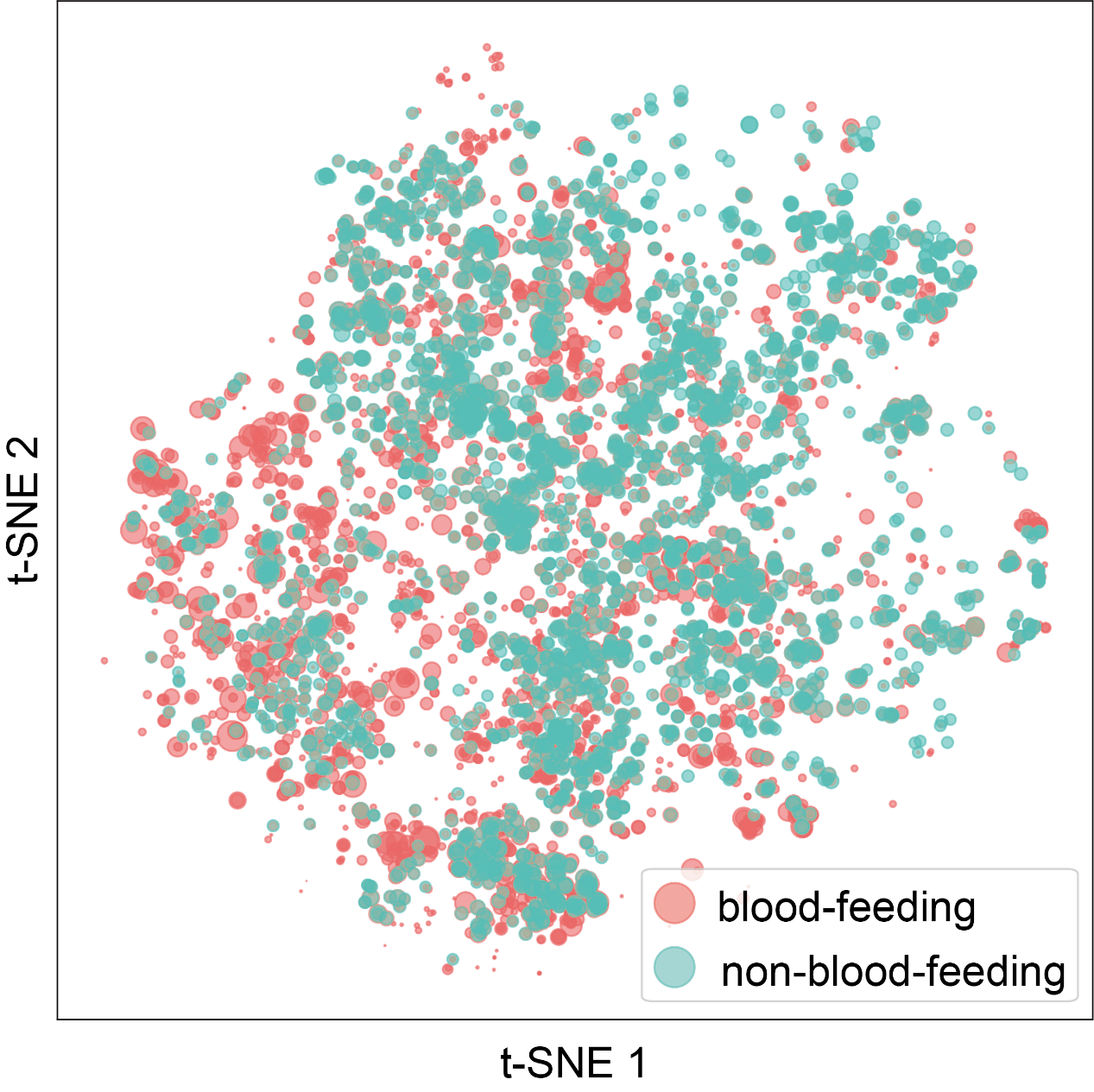

**Fig. S16 | The proportions of strong-binding VOCs for blood-feeding (red) and non-blood-feeding (green) mosquitoes in the 14,499 VOC dataset show substantial overlap**

### Performance of insectOlf on an independent larval *D. melanogaster* OR dataset

In addition to our experimentally tested dataset on *Bactrocera dorsalis*, we extracted an independent dataset from a published larval *D. melanogaster* OR study^10^ for further validation. This dataset comprises 608 OR–VOC response pairs derived from 19 ORs and 32 VOCs. For each OR–VOC pair, the response value at the highest tested concentration was taken as the experimental readout. OR sequences were retrieved from UniProt, and VOC structures were obtained from PubChem using CAS numbers provided in the original publication. Pairs with incomplete sequence or structural information were excluded from further analysis.

As shown in Fig. S17, the cumulative hit rate (CHR) curve (left) rises steadily above the diagonal throughout the ranking, indicating that positive hits are consistently enriched among top-ranked predictions relative to random expectation. The enrichment factor (EF) at the 20% cutoff was 1.48, corresponding to a 48% improvement over random selection. The AUROC and AUPRC values were 0.63 and 0.45, respectively, suggesting moderate discriminative power. Notably, this dataset involves a different experimental system (larval vs. adult OR expression), distinct OR-vOC combinations, and data from a different laboratory, all of which may contribute to variability in absolute performance metrics. Nonetheless, the overall enrichment above random expectation supports the generalizability of insectOlf across independent experimental contexts.

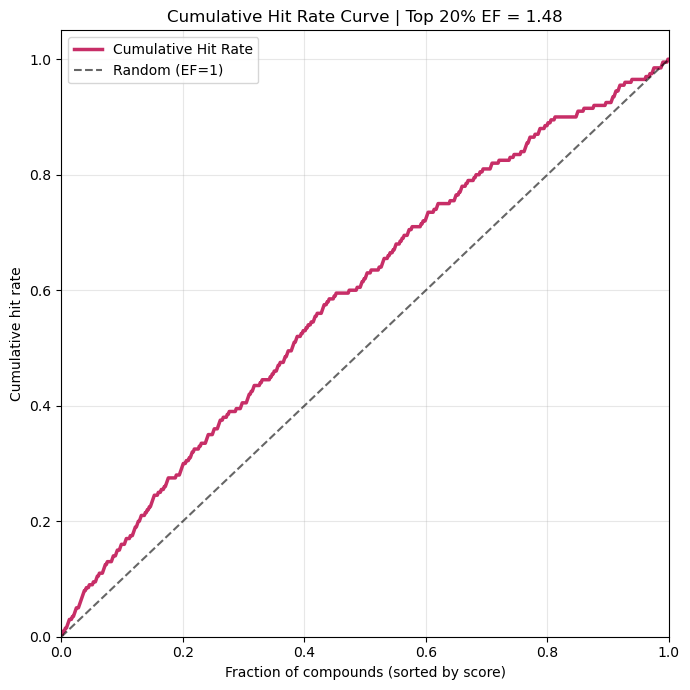

**Fig. S17 | Cumulative Hit Rate Curve of larval *D. melanogaster* OR**

1	Karpe, S. D., Tiwari, V. & Ramanathan, S. InsectOR—Webserver for sensitive identification of insect olfactory receptor genes from non-model genomes. *PLOS ONE* **16**, e0245324, doi:10.1371/journal.pone.0245324 (2021).

2	Li, Q. *et al.* iORbase: A database for the prediction of the structures and functions of insect olfactory receptors. *Insect Sci* **30**, 1245-1254, doi:10.1111/1744-7917.13162 (2023).

3	Holm, L. Using Dali for Protein Structure Comparison. *Methods in molecular biology (Clifton, N.J.)* **2112**, 29-42, doi:10.1007/978-1-0716-0270-6_3 (2020).

4	Hellwich, K.-H., Hartshorn, R. M., Yerin, A., Damhus, T. & Hutton, A. T. Brief guide to the nomenclature of organic chemistry (IUPAC Technical Report). **92**, 527-539, doi:doi:10.1515/pac-2019-0104 (2020).

5	Chi, H. *et al.* Genomic and phenotypic evidence support visual and olfactory shifts in primate evolution. *Nature Ecology & Evolution*, doi:10.1038/s41559-025-02651-5 (2025).

6	Lyu, J. *et al.* Ultra-large library docking for discovering new chemotypes. *Nature* **566**, 224-229, doi:10.1038/s41586-019-0917-9 (2019).

7	Wright, C. J., Stevens, L., Mackintosh, A., Lawniczak, M. & Blaxter, M. Comparative genomics reveals the dynamics of chromosome evolution in Lepidoptera. *Nature Ecology & Evolution* **8**, 777-790, doi:10.1038/s41559-024-02329-4 (2024).

8	Letunic, I. & Bork, P. Interactive Tree of Life (iTOL) v6: recent updates to the phylogenetic tree display and annotation tool. *Nucleic Acids Research* **52**, W78-W82, doi:10.1093/nar/gkae268 (2024).

9	Orme, D. *et al.* *caper: Comparative Analyses of Phylogenetics and Evolution in R*, <<https://CRAN.R-project.org/package=capers>> (2023).

10	Si, G. *et al.* Structured Odorant Response Patterns across a Complete Olfactory Receptor Neuron Population. *Neuron* **101**, 950-962.e957, doi:10.1016/j.neuron.2018.12.030 (2019).
